# AlphaFold-Driven Structural Proteomics Reveals Extensive Cellulosome Machinery in Human Ruminococcal Symbionts

**DOI:** 10.64898/2026.02.05.704116

**Authors:** Christine Minor, Allen Takayesu, Mark A. Arbing, Sung Min Ha, Robert P. Gunsalus, Matteo Pellegrini, Michael R. Sawaya, Robert T. Clubb

**Author notes:** To whom correspondence should be addressed: Dr. Robert T. Clubb, Department of Chemistry and Biochemistry, University of California, Los Angeles, 602 Boyer Hall, Los Angeles, CA 90095., Dr. Michael R. Sawaya, UCLA-DOE Institute of Genomics and Proteomics, University of California, Los Angeles, 205 Boyer Hall, Los Angeles, CA 90095, **Email:** and. C.M. and A.T. contributed equally to this work. R.T.C. and M.R.S. are co-corresponding authors.

## Abstract

Cellulosomes are large, surface-displayed enzyme complexes that enable anaerobic bacteria to degrade recalcitrant plant polysaccharides, yet cellulosome-expressing bacteria are thought to be rare in the human gut. Here we show that extensive sequence divergence obscures the detection of many ruminococcal cellulosomes by conventional sequence homology-based methods. Using proteome-scale AlphaFold2 structural predictions, we uncovered a substantially expanded set of cellulosome-producing *Ruminococcus* species, including six previously unrecognized human symbionts. Structure-based clustering identifies several novel cohesin families that retain conserved folds despite extreme sequence divergence and define distinct, phylogenetically conserved cellulosome architectures. The analysis reveals *R. callidus* and related human symbionts encode elaborate cellulosomes that are invisible to sequence-based annotation. Similarly, *R. difficilis*, a human gut symbiont, is found to produce an atypical cohesin-based assembly enriched in amylases and related starch-binding proteins that may enable this microbe to degrade resistant starches that evade digestion in the upper gastrointestinal tract. Together, these findings reveal that ruminococcal cellulosomes are far more prevalent and diverse than previously appreciated and demonstrate the power of structural proteomics to uncover deeply divergent functional systems in the gut microbiome.

**Significance Statement:** Plant cell wall polysaccharides are a major dietary carbon source, yet their degradation relies on rare, highly specialized microbial enzyme assemblies known as cellulosomes, which have long appeared uncommon in the human gut. Using proteome-scale structure prediction combined with experimental validation, we show that cellulosomes are far more widespread and structurally diverse in human-associated *Ruminococcus* species than previously appreciated. We identify multiple new cohesin families and reveal distinct cellulosome architectures likely adapted to degrade different dietary substrates. Together, these findings redefine the distribution and evolution of cellulosomes in gut microbes and demonstrate the power of structural proteomics to uncover deeply diverged biological systems.

## Introduction

Lignocellulosic plant biomass (LCB) is the most abundant source of organic carbon in the biosphere, yet its inherent resistance to degradation limits its utility as a nutrient and as a feedstock for renewable commodity production.^1^ Its cellulose, hemicellulose, and lignin components form a densely interlinked matrix, where the crystalline, hydrogen-bonded structure of cellulose restricts conversion to glucose monomers.^2,3^ To overcome this barrier, several anaerobic bacteria produce massive, multi-enzyme complexes known as cellulosomes, which concentrate glycoside hydrolase activity at the microbial surface to efficiently degrade cellulose, hemicellulose, resistant starch, and other complex dietary fibers.^4–7^ While cellulosome-producing bacteria are critical for degradation in the rumen of herbivores and the hindgut of non-ruminant animals, they are exceptionally rare in the human colon, with only five species identified among the ∼5,000 resident taxa. Investigation of cellulosomal components within ruminant-dwelling microbes reveals that the primary sequences of these components have diverged substantially, making them undetectable by conventional sequence-based approaches. However, we demonstrate that by using proteome-scale structural predictions, these complexes can be readily detected. We establish that cellulosomes are more widespread than previously appreciated, specifically in the human gut, which hosts as many as 11 distinct species capable of degrading diverse plant biomass.

Cellulosomes organize carbohydrate-active enzymes (CAZymes) at the microbial surface, enhancing lignocellulose degradation through coordinated activity and substrate targeting.^4–6^ These complexes are built around scaffoldin proteins, which contain cohesin domains that bind to dockerin-fused glycoside hydrolases (DocGHs) (**Fig. 1A**). Scaffoldins often contain carbohydrate-binding modules (CBMs) that direct the complex to cellulose, as well as dockerins that enable scaffoldin-scaffoldin interactions to increase the complexity, loading capacity, and range of the cellulosome. In the prototypical cellulosome from *Acetivibrio thermocellus* (also known as *Clostridium thermocellum*), the primary scaffoldin recruits diverse GHs — including endoglucanases, exoglucanases, and β-glucosidases — and attaches to cell wall–associated scaffoldins to form a highly organized enzymatic network.^8,9^ A core set of DocGHs (GH5, GH9, GH10, GH11, GH43, GH48) provide broad lignocellulose-degrading activity, supplemented by other CAZymes such as polysaccharide lyases, esterases, and oligosaccharidases.^6,7,10^ The spatial organization of these enzymes enhances synergy, improves substrate targeting, and promotes close interactions between the microbial surface and polysaccharide substrates, collectively maximizing degradation efficiency.^11–13^

**Figure 1.**
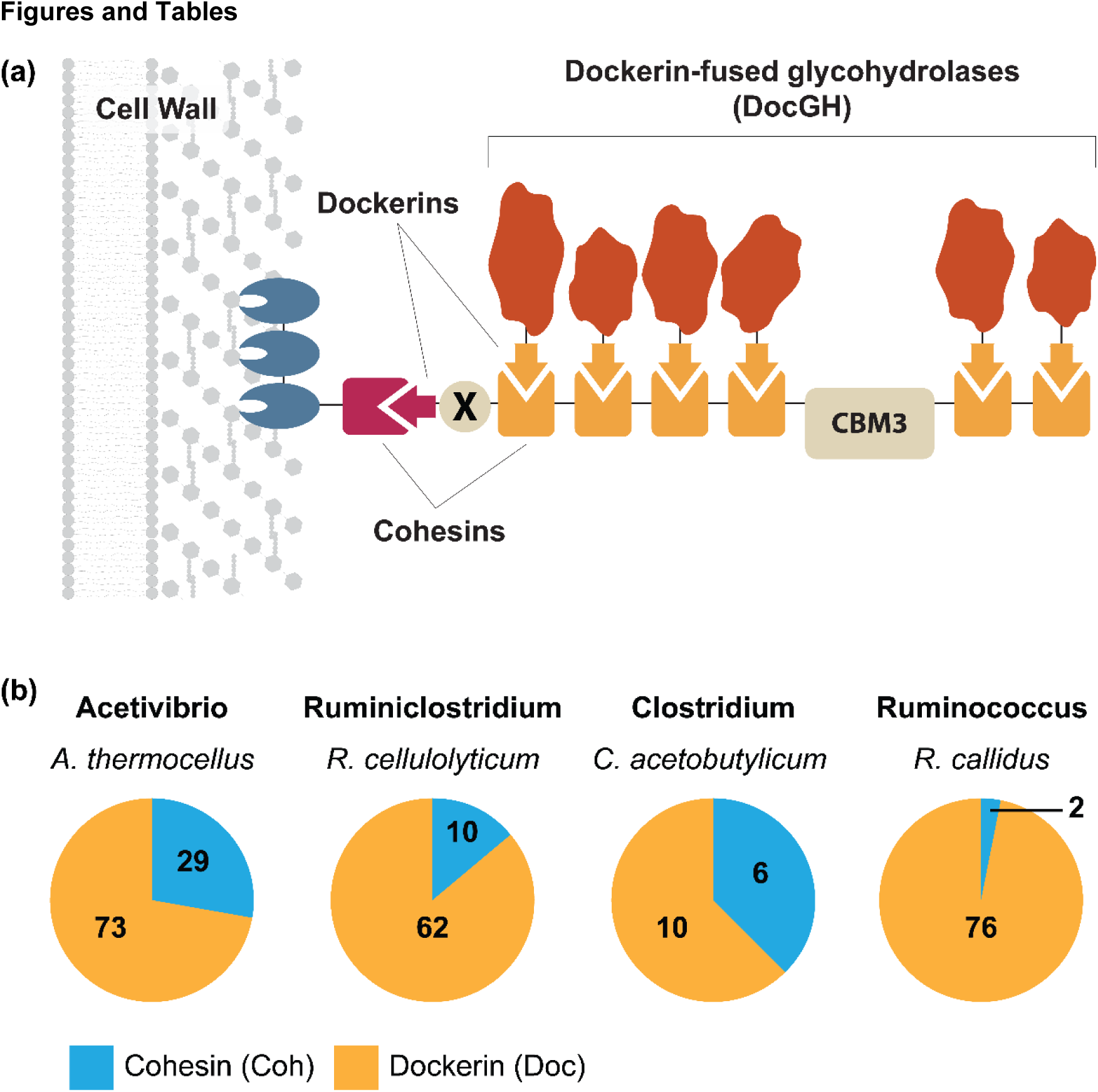
Cellulosomes in anaerobic bacteria. **(A)** Schematic showing a prototypical surface displayed cellulosome and cohesin-dockerin interactions. An anchoring scaffoldin associates with the cell surface and typically contains a type II (red) cohesin domain that interacts with a type II dockerin domain (red) located within a primary scaffoldin. Most primary scaffoldins also contain a series of type I cohesins (orange) that bind to dockerin-fused glycohydrolases (DocGHs) via a type I dockerin. In many instances the primary scaffoldin also contains a cellulose binding carbohydrate-binding module family 3 domain (CBM3). Most bacteria contain significantly more complex cellulosomes than shown here that are built from additional types of cohesin-dockerin interactions. **(B)** Cohesin and dockerin domain counts in representative cellulosome-producing bacteria from the *Acetivibrio* (*Acetivibrio thermocellus*), *Ruminoclostridium* (*Ruminoclostridium cellulolyticum*), *Clostridium* (*Clostridium acetobutylicum*), and *Ruminococcus* (*Ruminococcus callidus*) genera. Cohesin and dockerin counts shown are determined from InterProScan sequence profiles (Cohesins: type I – cd08548, type II – cd08547, type III – cd08759, cohesin domain – PF00963; Dockerins: type I – cd14256, type II – cd14254, type III – cd14255).

*Ruminococcus* species are key members of the gut and rumen microbiomes, with changes in their abundance linked to metabolic disorders, inflammatory bowel disease, and other gastrointestinal conditions.^14–16^ However, identifying cellulosomes in these microbes has been challenging using conventional homology-based methods, as the cohesin domains within their scaffoldins appear to have highly divergent sequences. For example, BLAST or Hidden Markov Model (HMM) searches of the human symbiont *R. callidus* detect only two cohesin domain-containing scaffoldins, despite the fact that its genome encodes 76 dockerin-fused enzymes (**Fig. 1B**).^17^ Other *Ruminococcus* species show similarly low scaffoldin counts, in contrast to richer repertoires in *Acetivibrio*, *Clostridium*, and *Ruminiclostridium*.^32^ Thus far, sequence-based methods have identified only five human gut cellulosome-producing *Ruminococcus* species (*R. champanellensis*, *R. bromii,* and recently identified candidate species: *R. hominiciens*, *R. primaciens,* and *R. ruminiciens*).^18^

In this study by employing proteome-scale AlphaFold predictions, we discovered six additional human symbionts that produce cellulosomes and revealed previously unrecognized scaffoldin-like domains across nearly all analyzed species, indicating more complex cellulosomes than previously appreciated.^19^ Structural clustering, validated by X-ray crystallography, identified novel cohesin families that uncovered phylogenetically conserved cellulosome architectures potentially tailored to degrade different types of biomass. These findings indicate that ruminococcal cellulosomes have been systematically underestimated and establish structural proteomics as a powerful approach to uncover deeply divergent functional systems in bacteria.

## Results

### Detection of novel cohesin-containing cellulosomes in ruminococcal bacteria

Reasoning that scaffoldin sequences in *Ruminococcus* spp. are highly divergent, we applied AlphaFold2 (AF2) at a proteome-scale to uncover their cohesin domains (**Fig. 2A**). Initially, 124 ruminococcal genomes in the NCBI taxonomy database (Accessed: August 2025) were translated and their primary sequences searched with InterProScan to identify dockerin domain-containing proteins using established HMM profiles within Pfam and the conserved domain database (CDD).^20,21^ These genomes contain all documented *Ruminococcus* species in the List of Prokaryotic Names with Standing in Nomenclature (LPSN), including several that have subsequently been reclassified as *Blautia, Mediterraneibacter, Trichococcus,* or *Hominimerdicola* species.^22–24^ A total of 56 of the 114 genomes contain ≥15 dockerin-encoding genes compatible with the presence of a cellulosome, of which 42 genomes were carried forward for further analysis as they correspond to non-redundant species as defined by ProGenomes (**Table S1**). To identify cohesin-like modules, a locally implemented version AF2 was used to predict their protein structures (41,171 atomic structure predictions across 42 proteomes). The structures were parsed into their component domains using UniDoc, and structural similarity to known cohesin structures assessed by calculating a TM-score; a metric ranging from 0 (no similarity) to 1 (identical folds) that quantifies their structure similarity.^25,26^ Domains were classified as a cohesin if their TM-score exceeded a threshold value of 0.65 when compared to at least one of three representative cohesin structures (type I (PDB 1OHZ), type II (PDB 1TYJ), and type III (PDB 2ZF9) cohesins). This threshold reliably separates high-from low-confidence cohesin hits. For example, 2,870 protein structures were predicted for *R. callidus* VPI 57-31 (ATCC 27760), which UniDoc parsed into 6,164 domains. Comparing these to a representative type I cohesin (PDB: 1OHZ) identified 17 hits with TM-scores ≥0.65 (red in **Fig. 2B**), which also have uniformly low root mean square differences (RMSDs) to the reference (red in **Fig. 2C**). Similar trends are observed for *R. champanellensis* and *R. flavefaciens* predictions, further supporting the threshold selections (**Fig. S1**).

**Figure 2.**
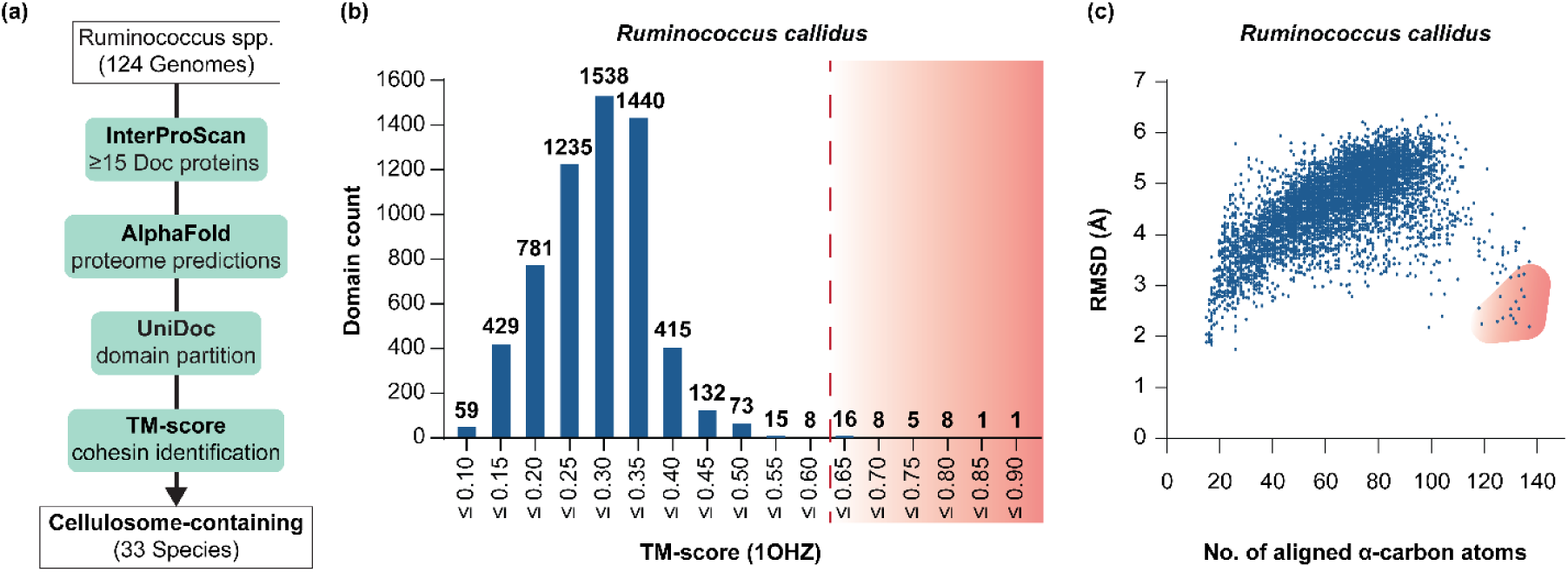
AlphaFold-based screen to identify novel scaffoldins. **(A)** Strategy used to identify cohesin domains in *Ruminococcus* genomes. A total of 124 *Ruminococcus* genomes and metagenome-assembled genomes (MAGs) were initially analyzed by InterProScan. Of these, 42 genomes were carried forward for structure prediction using AF2, parsed into structured domains using UniDoc, and inspected for cohesin-like modules using TM-align. **(B)** Representative plot showing the AF2 predicted structures within the *R. callidus* proteome, binned according to their calculated TM-score; 6,164 individual domains were compared to a type I cohesin (PDB: 1OHZ). The threshold TM-score used for cohesin domain identification is shown as a dashed red line that separates a large low-scoring population of unrelated structures (left) from a high-scoring structurally related population (right). **(C)** Scatter plot of RMSD α-carbon coordinate difference versus number of aligned α-carbon atoms for the data shown in panel (B). The plot shows that domains with high TM-scores (shaded red) exhibit small coordinate deviations with the representative type I structure, highlighting robustness of the TM-score as a discriminator. Similar trends are observed when the *R. flavefaciens* and *R. champanellensis* proteomes are analyzed (**Fig. S1**).

Based on AF2 structural proteomics 33 of the 42 bacterial genomes contain genes encoding cellulosomes (their predicted proteomes contain ≥15 dockerin-containing proteins and at least one potential scaffoldin that harbors ≥2 and ≥1 cohesin and dockerin modules, respectively) (**Fig. 3**). Of these, nine cellulosome-displaying species were only detectable using AF2 and otherwise evade sequence homology-based methods, including several human symbionts (discussed below). Moreover, in all but one of the 33 cellulosome-displaying microbes, AF2 detected novel cohesin-containing scaffoldins that were otherwise invisible to sequence-homology based methods (the lone exception being *R. bromii*). In several instances, AF2 structural searches are more than double the number of cohesins detected in each species (e.g. in *R. callidus* and *R. difficilis*) (**Fig. S2A**). Specifically, a total of 538 cohesin domains were detected using AF2, BLAST or InterPro HMM profiles, of which 177 were only identified by AF2 (hereafter called AF2-only cohesins) (**Fig. S2B**). Interestingly, a sequence profile designed to detect β-sandwich folds that resemble cohesins is more effective in identifying ruminococcal cohesins (CATH-GENE3D profile G3DSA:2.60.40.680). However, searches with this profile lead to false-positive results, as they also capture non-cohesin proteins that adopt similar protein folds with distinct functions (e.g. carbohydrate binding modules).^27,28^ Cohesin identification by structure is robust, since, with only a few exceptions, all cohesins detected based on their primary sequence are also identified using AF2 (**Fig. S2B**). Similar gains are not observed when representative cellulosome-producing bacteria from other genera (*Clostridium, Acetivibrio*, and *Ruminiclostridium*) are searched for cohesins using AF2, suggesting that their cohesins are less divergent and thus detectable using established sequence-homology based methods (**Fig. S2A**). None of their genomes encoded cellulosome components suggesting that this adaptation is a distinct feature of ruminococcal bacteria.^29^ In addition to revealing a new phylogenetic clade of cellulosome producing Ruminococcus species, these results underscore the power of AF2-based structural proteomics to complement sequence based homology methods in expanding the cohesin space resulting in an architecturally richer set of ruminococcal cellulosomes than previously appreciated.

**Figure 3.**
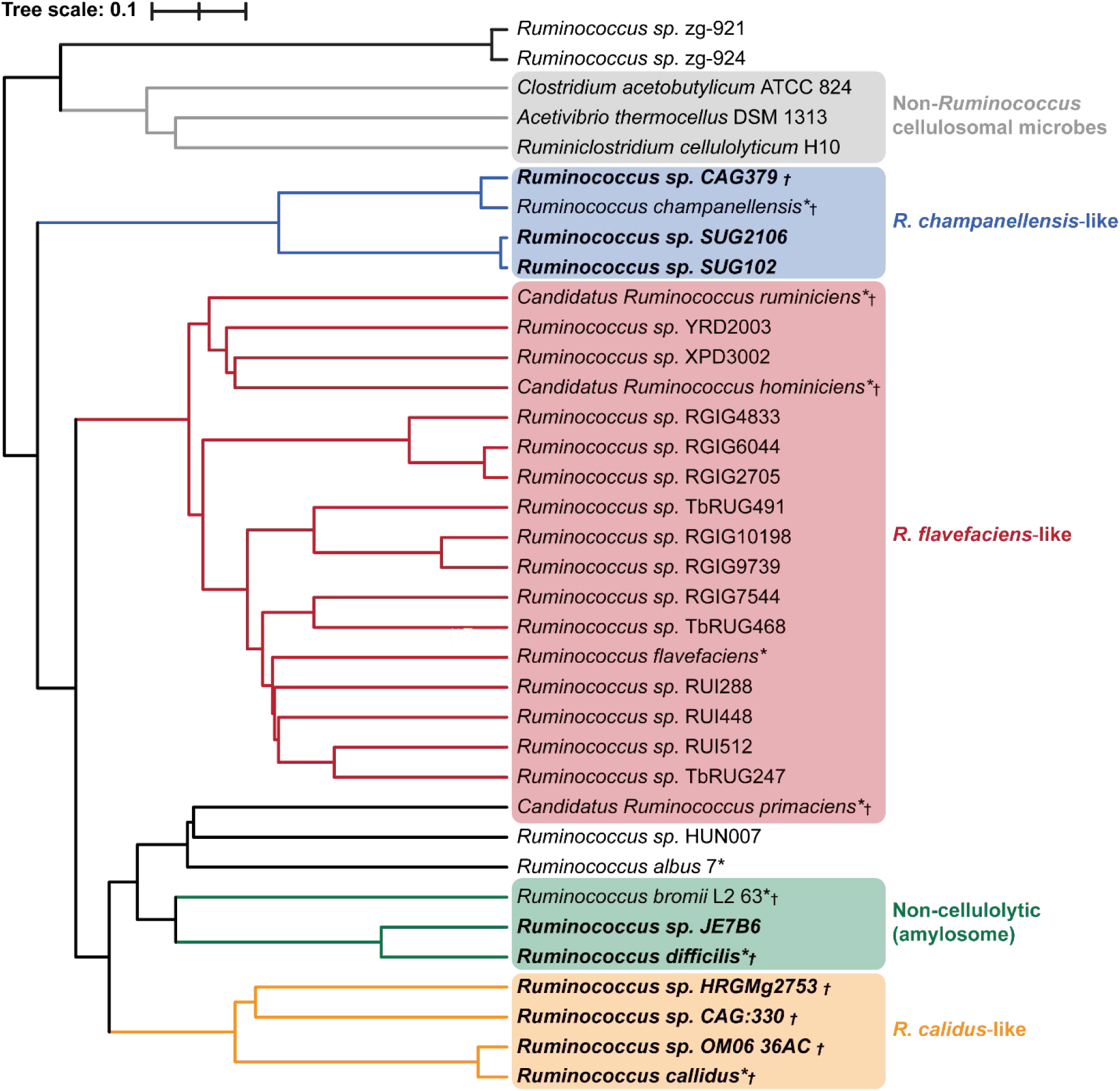
Phylogeny of cellulosome-producing *Ruminococcus* species. Phylogenetic tree of 33 cellulosome-producing *Ruminococcus* species based on their whole-genome Average Nucleotide Identity (ANI). Species are colored based on their cellulosome archetype: *R. flavefaciens*-(red), *R. champanellensis*-(blue), and *R. callidus*-like (gold) cellulosomes in addition to three non-cellulolytic or amylosomal species (green). Also shown are distantly related cellulosome-displaying bacteria from other genera, demonstrating their sequence divergence (gray, *Acetivibrio*, *Clostridium*, and *Ruminiclostridium* genera). *Ruminococcus* species designated by the LPSN are annotated with an asterisk (*) and those that are human-associated are annotated with a dagger (†). Species that are bolded were discovered in this study as cellulosome-producing bacteria. The tree was constructed using FastANI and clustered using the UPGMA algorithm.

### Cohesins in ruminococcal bacteria adopt non-canonical structures with diverse primary sequences

To gain insight into their roles in cellulosome assembly, we systematically compared the structures of AF2-only detected cohesins by constructing a dendrogram based on their pairwise TM-scores (**Fig. 4**) that fall into seven distinct groups. This analysis revealed a wide range of diversity, including the presence of several structural cohesin groups (groups 4 to 7) whose members share TM-scores ≥0.75. Their predicted structures are novel, differing from cohesins in the protein databank (PDB) that are identifiable based on their primary sequences; HMM profiles classify many of the cohesins in the PDB as type I to III family members, which based on their structures form structural groups 1 to 3, respectively (**Fig. S3**). Notably, ruminococci contain only a few cohesins with structures or primary sequences that are homologous to type II cohesins. which in other bacterial genera are in cell wall associated scaffoldins that tether cellulosomes to the microbial surface. Instead, in ruminococci, this function appears to be performed by a range of sequence-divergent modules originating from groups 1, 3-4, and 7 (**Figs. 6, S7**). Interestingly, the structures of several of the AF2-only cohesins resemble canonical group 1 cohesins, even though their primary sequences do not conform to the type I HMM profile. This sequence-structure discordance is consistent with a sequence similarity network (SSN) analysis of the AF2-only detected cohesins, as members of each structural group can be clustered into several, distinct families of proteins that share >30% sequence identity (**Fig. S4**). Despite the significant sequence divergence, the data suggest that ruminococci leverage cohesin structural conservation to generate scaffoldins needed to construct more elaborate cellulosome architectures.

**Figure 4.**
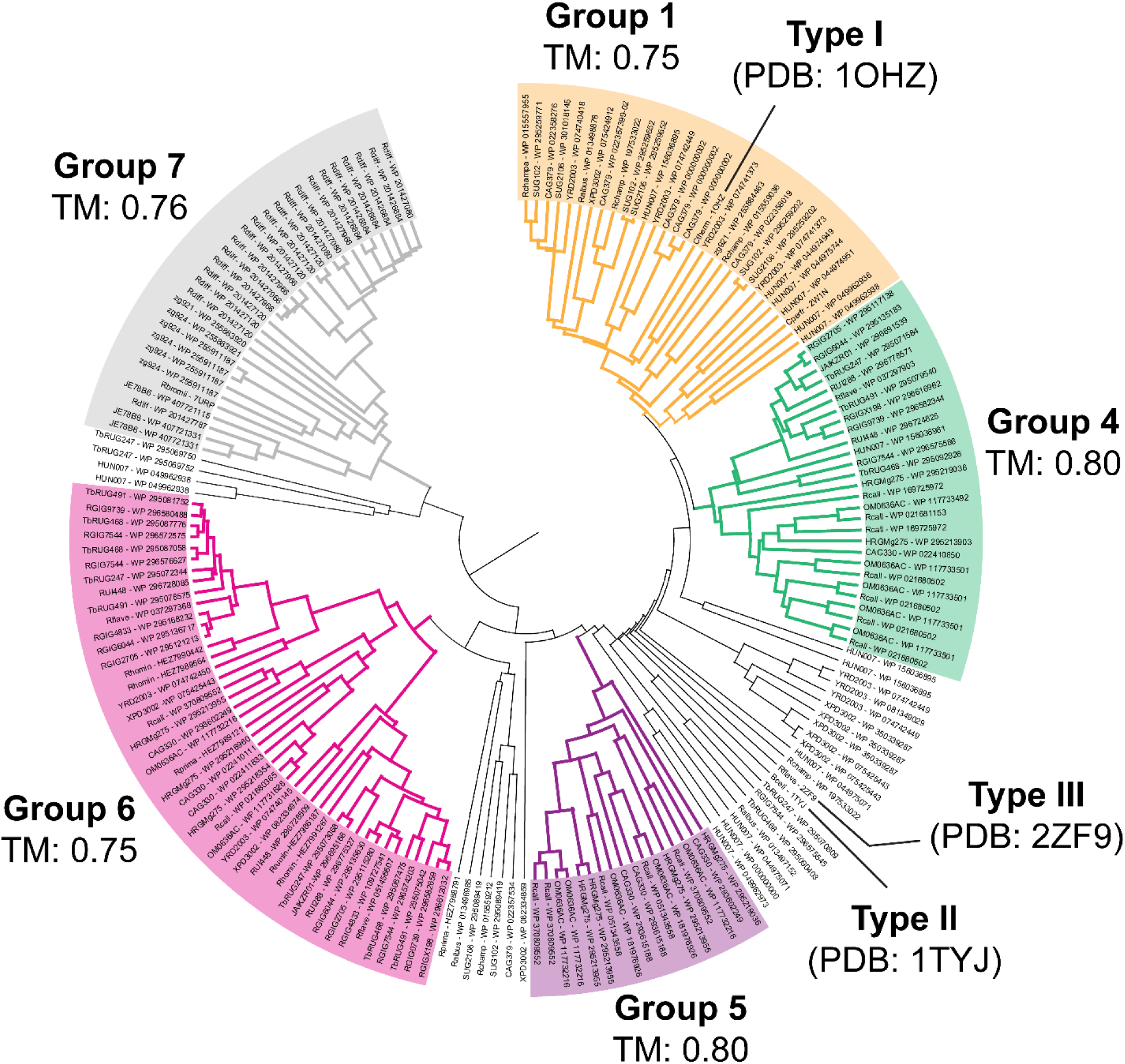
Structure-based dendrogram of AF2-only detected ruminococcal cohesins. Structural dendrogram of the 177 *Ruminococcus* cohesins that could only be detected with AF2. The dendrogram was constructed from an all-on-all TM-score matrix, with proteins sharing TM-scores ≥0.75 clustered into groups 1, and 4-7. For reference, the dendrogram also shows three representative type I to III cohesin structures (PDB 1OHZ, 1TYJ, and 2ZF9) that form structural groups 1 to 3, respectively (**Fig. S3**).

**Figure 5.**
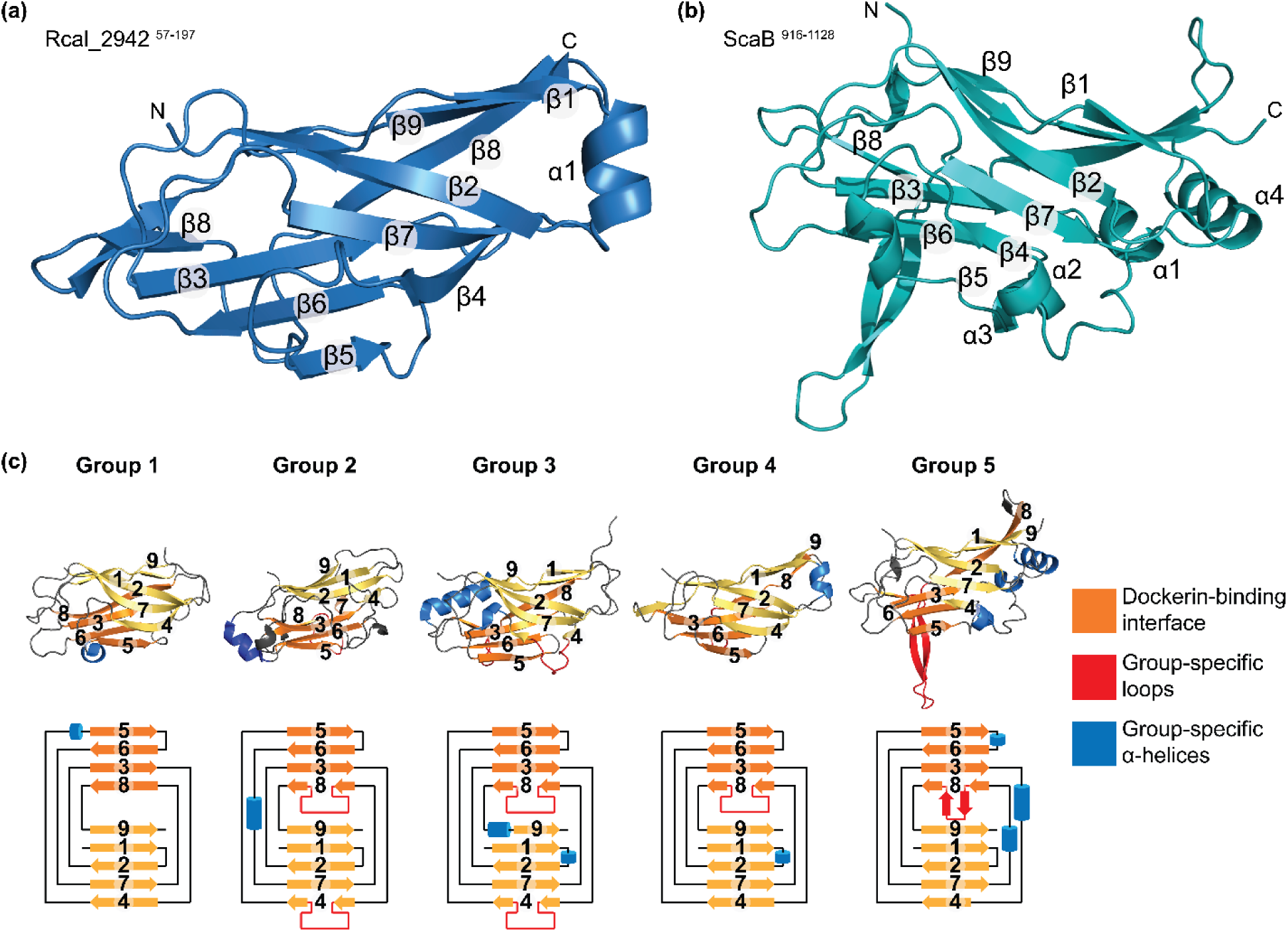
Structurally novel cohesins in ruminococcal bacteria. Cartoon representation of representative crystal structures **(A)** Group 4 (Rcal_2942, S57-V197) and **(B)** Group 5 (Rcal_0153, Q916-P1128) cohesins. **(C)** Secondary structural topology diagrams comparing different types of cohesins. Representative atomic structures from the PDB are shown for type I (group 1, PDB: 1OHZ), type II (group 2, PDB: 5G5D), and type III (group 3, PDB: 2ZF9) sequence families. Topologies determined for crystal structures reported in this study for group 4 (Rcal_2938, K547-P685) and group 5 (Rcal_0153, Q916-P1128) proteins are also shown. Cohesin structures contain a nine-stranded β-sandwich jelly roll with strands 5, 6, 3, and 8 (orange) forming the dockerin-cohesin interface. Group-specific accessory helices (blue) and loops (red) are highlighted.

**Figure 6.**
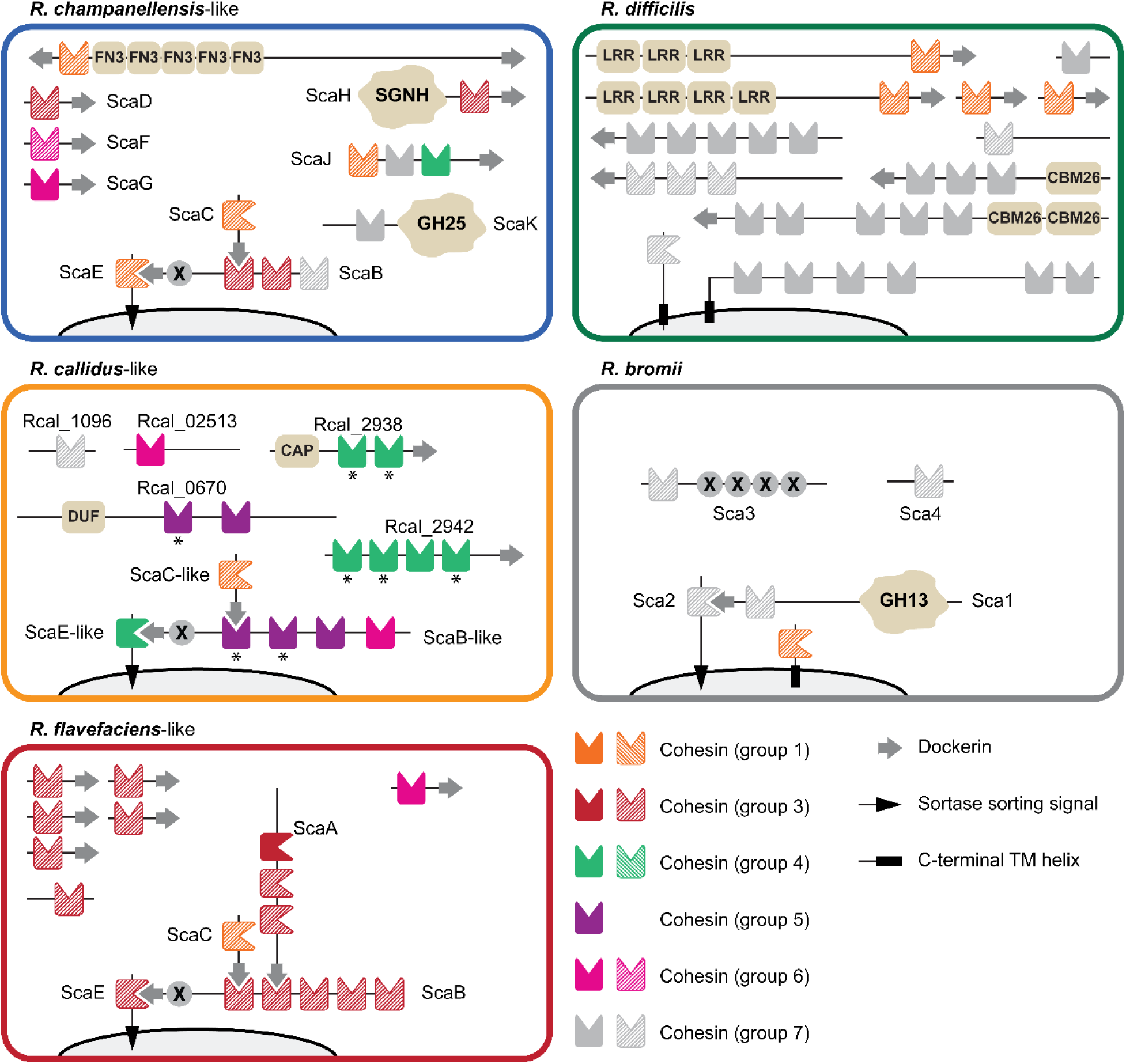
Cellulosome archetypes in human symbionts. Cartoon schematic of the scaffoldins in human-associated bacteria. Cohesins that can only be detected with AF2 have solid shading, whereas cohesins that can also be detected using sequence homology-based methods are hatched. Additional domains within the scaffoldins are as follows: **FN3**, Fibronectin Type III; **X**, X2 domain; **SGNH**, SGNH Hydrolase; **GH**, glycoside hydrolase; **CAP**, Cysteine-rich secretory / Antigen 5 / Pathogenesis-related 1 domain; **DUF**, Domain of Unknown Function (DUF5620), **LRR** – Leucine Rich Repeat, **CBM26** – Carbohydrate Binding Module Family 26. The *R. callidus-* and *R. difficilis*-like cellulosomes could only be detected using AF2.

To confirm the presence of novel cohesins in ruminococcal bacteria, X-ray crystallography was used to resolve the structures of eight predicted group 4 and 5 cohesins from *R. callidus* (marked with an asterisk, **Fig. 6**). Structures were determined at resolutions ranging from 1.2-2.0 Å and 1.8-2.8 Å, for five group 4 (coh1 and coh2 from Rcal_2938; coh1, coh2, and coh4 from Rcal_2942) and three group 5 (coh3 and coh4 from Rcal_0153, coh1 from Rcal_0670) cohesins, respectively (**Table S2**). Overall, the experimentally determined and predicted structures are in good agreement, and exhibit backbone coordinate RMSD values ranging from 0.33-1.03 Å, with the maximum coordinate deviations occurring in their loops and termini (**Fig. S5**).

Despite lacking significant sequence homology, group 4 and 5 structures adopt a conserved cohesin-like fold that is constructed from a two-sheeted β-sandwich jellyroll that contains 9 antiparallel β-strands (**Fig. 5A, B**). While a detailed analysis of their differences with canonical cohesin structures is beyond the scope of this manuscript, a comparison reveals additional α-helices and/or loops distinguish group 4 and 5 proteins from other types of cohesins (**Fig. 5C**). The group 4 cohesins determined by crystallography show only a few deviations from the canonical cohesin fold. Notably, four of the five experimentally determined group 4 cohesins contain an additional α-helix inserted between β-strands 5 and 6 that was not predicted by AF2, underscoring the importance of experimental structural validation (**Fig. 5A**). Group 5 proteins exhibit more significant differences with canonical cohesins, as they contain an elongated β-hairpin that is inserted into strand β8 (red, **Fig. 5B, C**). The ∼20 residue β-hairpin is a large deviation from the smaller ∼9-13 residue β-flap insertions within the β8 strand commonly seen in group 2 (e.g. PDB: 5LXV), 3 (e.g. PDB: 1TYJ, 4UYP), and 4 cohesins.^30–32^ As with these β8 flaps, the β-hairpin arm within group 5 cohesins resides near the presumed cohesin:dockerin interface, suggesting it may play a role in forming specific contacts for dockerin binding. Indeed, in all three group 5 cohesins determined via x-ray crystallography, the extended β-hairpin arms contained several exposed hydrophobic residues to potentially mediate specific non-covalent cohesin:dockerin binding. Experimentally determined crystal structures of group 5 cohesins also contain accessory α-helices that decorate the cohesin core (group 5, blue, **Fig. 5C**), which may form an additional protein interaction surface. Taken together, our findings show that the structures and sequences of ruminococcal cohesins extend well beyond canonical types I–III, forming diverse structural groups with unique fold embellishments that likely confer dockerin domain binding specificity.

### Structural classification reveals distinct ruminococcal cellulosome architectures

Using our AF2-based classification scheme, we analyzed the scaffoldin content of cellulosomes in ruminococci species to gain insight into their architecture and functions in LCB degradation. Most cellulosome-producing ruminococcal species contain phylogenetically conserved cellulosomes with architectures resembling those found in either *R. flavefaciens, R. champanellensis*, or *R. callidus* (**Fig. 3**). Microbes displaying *R. callidus*-type cellulosomes exclusively populate the human gut and are only detectable using our AF2-based method. Conversely, bacteria displaying *R. flavefaciens*– and *R. champanellensis*-like structures reside within either the human or herbivore intestinal tracts and have cellulosomes with components that can generally be detected using conventional sequence-homology based methods.

The largest number of ruminococcal species characterized to date are phylogenetically related to the rumen bacterium *R. flavefaciens* and produce architecturally related cellulosomes with scaffoldins containing type III cohesins that are identifiable based on their primary sequence (hatched cohesins in **Fig. 6**).^17^ These microbes possess a core set of scaffoldins (ScaA, ScaB, ScaC and ScaE) that are thought to associate with one another via non-covalent cohesin-dockerin interactions and to be collectively attached to the cell surface via the ScaE scaffoldin, which is covalently attached to the peptidoglycan.^17,33^ These cellulosomes also contain ∼5-6 monovalent adaptors that may promote type-switching between cohesin-dockerin pairs and/or the assembly of fiber-like structures that extend from the cellulosome.^7,34^ Typically the bacteria with *flavefaciens*-like cellulosomes inhabit the rumen and possess a strong arsenal of DocGH proteins with activities primarily against cellulose (GH5, 9, 48) and hemicellulose (GH10, 11, 26, 43) (**Table S3**). Additional GH families with more specialized roles such as GH97 (oligosaccharidases) and GH14 (amylases) are limited but nonetheless complement the breakdown of diverse glycosidic bonds. *R. champanellensis*-like cellulosomes contain core scaffoldins resembling the *R. flavefaciens* archetype that are generally identifiable based on their primary sequences (**Fig. 6**). However, in these microbes the core is elaborated with a distinct array of dockerin-fused cohesins, including SGNH-fused ScaH-like and GH25-fused ScaK-like scaffolding,^35^ as a well as a massive ∼2,800 residue protein that contains a single cohesin domain, several Fibronectin Type-II (FN3) motif repeats, and both N– and C-terminal dockerin modules. These structures are found in both human and herbivore-associated bacteria and harbor an enzyme complement that may facilitate the digestion of a diverse array of carbohydrates; they contain up to 57 DocGH genes that presumably degrade cellulose (GH5, 9) and hemicellulose (GH43), but also a diverse set of enzymes with auxiliary functions as classified by the CAZyme database (**Table S3**).

Surprisingly, the AF2 predictions reveal that *R. callidus* and closely related human symbiotic species produce a third, architecturally unique cellulosome that is only detected when structure homology-based methods are employed (cohesins with solid shading in **Fig. 6**). In these microbes, AF2-based methods uncovered an elaborate cellulosome formed by eight scaffoldins that harbor novel group 4, 5, and 6 cohesins. Although they contain sequence divergent cohesins, three proteins resemble ScaB-, ScaC-, and ScaE-like core scaffoldins based on their overall domain arrangement (only the ScaC-like protein was detectable based on its primary sequence).^17^ Moreover, as in *flavefaciens*– and *champanellensis*-like cellulosomes, these core scaffoldins may be tethered to the cell surface by interactions with a ScaE-like anchoring scaffoldin covalently anchored to the peptidoglycan via an LPXTG motif. This apparent functional similarity between the three major cellulosome archetypes is further supported by the positioning of the genes encoding the ScaB– and ScaC-like proteins, which are located within a cluster resembling the *sca* gene cluster found in *R. champanellensis* and *R. flavefaciens*.^36^ In *callidus*-like cellulosomes the core structure is uniquely elaborated by five additional scaffoldins, including the two polyvalent adaptor scaffoldins, Rcal_2938 and Rcal_2942 that contain group 4 predicted cohesins. *R. callidus*-like cellulosomes are the only archetype that contain group 5 cohesins, all of which appear in the ScaB-like primary scaffoldin or as a cohesin dyad in Rcal_0670. A complete list of the scaffoldin components in *R. callidus* is provided in **Table S4**.

Nine human symbionts have the genetic capacity to produce cellulolytic cellulosomes (**Fig. 3**). These include *R. champanellensis and R. sp. CAG379* species that produce *champanellensis-*like structures, and *R. callidus, R. sp.* HRGMg2783, *R. sp.* CAG330, and *R. sp.* OMO636AC that produce *callidus*-like structures. In addition, recently sequenced metagenome-assembled genomes (MAGs) from candidate human associated *Ruminococcus* species *R. primaciens, R. hominiciens*, and *R. ruminiciens* possess *flavefaciens*-like cellulosomes that are also prevalent in rumen bacteria.^18^ Interestingly, the cellulosomes in each of these microbes likely confer cellulolytic activity as their genomes contain a significant number of LCB degrading DocGH enzymes. However, as compared to herbivore symbionts displaying related cellulosomes, these human symbionts are presumably less cellulolytic as their genomes encode a significantly lower number of DocGHs. For example, even though *R. hominiciens* possess a flavefaciens-like cellulosome, its genome encodes only 21 DocGHs versus 26 in the rumen bacterium *R. flavefaciens* (**Table S3**).

AF2 driven discovery also uncovered a novel cellulosome-like structure in the human symbiont *R. difficilis* which may degrade starch and related carbohydrates. This microbe produces an atypical cellulosome that is presumably directly attached to the microbial membrane via C-terminal transmembrane helices located in two anchoring scaffoldins, and contains a large number of scaffoldins with novel group 7 cohesins. Unlike cellulolytic cellulosome-producing *Ruminococcus* species, only a few genes encoding DocGH enzymes are present in *R. difficilis*’ genome; of its 80 dockerin domain-containing proteins, *R. difficilis* contains only a single DocGH with hemicellulase activity (GH53) and a handful of mannosidase domains. Instead, *R. difficilis* may specialize in degrading resistant starch that escapes digestion in the small intestine as its genome encodes for seven amylase-dockerin fusion proteins and two multi-cohesin scaffoldins that contain CBM26 modules known to bind starch products that may aid in substrate targeting.^37^ Indeed, *R. difficilis* is phylogenetically related to *R. bromii* (**Fig. 3**), a keystone human gut symbiont that specializes in degrading resistant starch which also contains an atypical cellulosome-like structure that houses amylase enzymes (**Fig. 6**).^38,39^ However, the cellulosomes in these microbes have diverged significantly at the primary sequence level, as only *R. bromii’s* cellulosome is detectable based on its primary sequence. Moreover, *R. difficilis’* cellulosome is unique, as several of its scaffoldins harbor binding domains that may promote microbial adhesion (Ig-like, LRRs, and mucin binding protein domains). A complete list of the scaffoldin components in *R. difficilis* is provided in **Table S5**. Collectively, these findings uncover multiple conserved yet previously hidden classes of ruminococcal cellulosomes and show that many human-associated species deploy structurally distinct architectures likely adapted to specific dietary polysaccharides.

Several previously unrecognized herbivore-associated microbes were also discovered to display cellulosomes with distinct architectures. For example, the rumen bacterium *R. sp.* HUN007 contains a cellulosome archetype in which the canonical core structure is replaced with monovalent and polyvalent adaptor scaffoldins that house group 4 and 7 cohesins, as well as a single group 4 cohesin that is presumably attached to the lipid bilayer via a transmembrane helix (**Fig. S7**). Its genome encodes a large number of DocGHs that presumably degrade LCB (**Table S3**), suggesting this microbe may be a significant contributor to cellulolytic activity in the rumen microbiota. Other rumen associated microbes of note include *R. JE7B6, R. zg921,* and *R. zg924* which have at least three transmembrane anchored singular cohesin proteins from groups 1, 4, and 7. A majority of their multi-cohesin containing proteins are from group 7, which is consistent with *R. bromii’s* scaffoldins. *R. JE7B6* contains a unique scaffoldin with an N-terminal α-amylase and C-terminal CBM26 which supports its dockerin profile of 6 dockerin-fused α-amylase proteins. Similarly, *R. zg921* and *R. zg924* contain four dockerin-fused α-amylase proteins and five dockerin-fused CBM26 modules (**Table S3**).

## Discussion

In this study, we show that cellulosomes in *Ruminococcus* species inhabiting the intestinal tracts of humans and other animals have been systematically underestimated due to the extreme sequence divergence of their cohesin domains. In these microbes, sequence-based HMM profiles uncovered a large number of DocGH encoding genes that are hallmarks of cellulosome-producing bacteria, but relatively few cohesin-containing scaffoldins to which these enzymes presumably bind (**Fig. 1B**). Reasoning that their primary sequences had diverged significantly to render them undetectable using conventional sequence homology-based methods, we used AF2 to predict 41,171 protein structures across 42 ruminococcal genomes and MAGS to search for cohesin-like modules capable of binding DocGH enzymes (**Fig. 2A**). By applying proteome-scale AF2–based structural analysis, we identify multiple classes of cohesin-containing scaffoldins that evade detection by conventional sequence-homology methods (**Fig. S2**), substantially revising current estimates of cellulosome distribution and complexity in gut-associated *Ruminococcus* species (**Fig. 3**). In particular, our results indicate that the apparent rarity of cellulosome-producing ruminococci in the human colon largely reflects the limitations of primary sequence-based detection methods rather than true biological absence. Instead, structure-based discovery reveals that cellulosomes are widespread among ruminococci, with many species possessing phylogenetically conserved architectures, including cellulosome systems that are selectively associated with human intestinal lineages.

A central insight from this work is that cohesin identity and function are constrained at the structural level rather than primary sequence. AF2-driven discovery uncovered at least four new cohesin groups (4–7) (**Fig. 4**), with representative group 4 and 5 cohesins from *R. callidus* confirmed by X-ray crystallography (**Fig. 5**). These novel cohesins are only detectable based on their structures and possess highly varied primary sequences (**Fig. S4**) that are distinct from traditional type I–III proteins whose structures have been previously documented in the PDB. They each retain the characteristic cohesin-like jelly-roll architecture, but also incorporate group-specific structural embellishments near the predicted dockerin-binding interface (**Fig. 5B**). In general, cohesin-dockerin interactions have been shown to be both species– and type-specific (e.g. only type I cohesins and type I dockerins produced by the same species interact with each other).^40,41^ It is widely accepted that dockerins have multiple binding modalities to cohesin domains that increase flexibility of the cellulsoome to conform to its target carbohydrate and here, the novel cohesin groups defined may contribute directly to cellulosome adaptability, potentially through unique dockerin-binding modes that are yet to be characterized.^42,43^ This suggests that the cohesin diversification predicted by AF2 may enable orthogonal cohesin-dockerin interaction networks within ruminococci, permitting the assembly of complex, hierarchical cellulosomes as has been observed in other genera.^33^

At the systems level, structural classification reveals that most ruminococcal cellulosomes segregate into a limited number of conserved architectural archetypes, exemplified by *R. flavefaciens*-, *R. champanellensis*-, and *R. callidus*-like systems (**Figs. 3,6**). These archetypes are defined not only by cohesin fold families, but also by the number and type of accessory scaffoldins that are present. *Flavefaciens*– and *champanellensis*-like architectures are broadly distributed across both herbivore and human intestinal tracts and are partially detectable using sequence homology. In contrast, *callidus*-like systems appear restricted to human-associated species and are entirely dependent on structure-based detection, underscoring the extent to which sequence divergence has obscured their recognition. Despite profound divergence at the amino acid level, these distinct architectures preserve a conserved organizational logic. In particular, they all encode scaffoldins that are structurally analogous to canonical ScaB-, ScaC-, and ScaE-type proteins, retain sca-like gene clustering, and are predicted to anchor to the cell surface via the covalent attachment of ScaE protein to the peptidoglycan (**Fig. 6**). This convergence suggests selective pressure to maintain overall cellulosome topology and spatial organization, even as individual cohesin and dockerin sequences diversify extensively. A total of nine human symbionts display one of these three architectures (**Fig. 3**). However, these ruminococci generally encode fewer DocGHs than their herbivore-associated counterparts, implying reduced cellulolytic potential despite retaining structurally elaborate cellulosomes (**Fig. S6**). This pattern likely reflects differences in substrate availability between the rumen and the human colon, where plant cell wall material is more limited, chemically heterogeneous, and often partially processed prior to microbial fermentation. Future work integrating structural proteomics with biochemical interaction assays, *in situ* expression studies, and ecological analyses will be required to define the surface architectures and physiological roles of these newly identified cellulosomes.

Our analysis also uncovers an atypical, starch-oriented cellulosome-like system in the human symbiont *R. difficilis*, highlighting the broader diversity of cohesin-based assemblies beyond classical cellulolytic paradigms (**Fig. 6**). The predominance of group 7 cohesins, the presence of a transmembrane-anchored scaffoldin, and the enrichment of amylase–dockerin fusion proteins suggest a system optimized for the degradation of resistant starch rather than lignocellulose. The phylogenetic proximity of *R. difficilis* to *R. bromii*, a well-established keystone starch degrader in the human gut, further supports the existence of a distinct amylosomal lineage that has diverged substantially at the sequence level while retaining a recognizable structural logic (**Fig. 3**). Collectively, our findings demonstrate that the number of *Ruminococcus* cellulosome producers rival those found in other genera and significantly expand the structural and sequence space of cohesin domains within cellulosomes. To our knowledge, a proteome wide application of AF2 to identify new protein families has not been applied, although intermediate scale studies have annotated select proteins using AF2 and Foldseek.^44,45^ More broadly, this work establishes structure-guided proteomics as a powerful framework for uncovering functional assemblies that lie beyond the reach of sequence-based annotation and for refining our understanding of how modular enzyme systems evolve, specialize, and persist within complex microbial ecosystems.

## Materials and Methods

### Sequence– and structure-based identification of cohesins in *Ruminococcus* species

Initially, 124 *Ruminococcus* genomes were accessed from the NCBI taxonomy database (Accessed: August 2025), consisting of 117 curated Reference Sequence (RefSeq) genomes obtained from the database and seven genomes from the Genbank database. To ensure we were looking at the most up to date list of *Ruminococcus* species, we compared this list to the *Ruminococcus* List of Prokaryotic names with Standing in Nomenclature (LPSN). The LPSN contained 32 reference genomes, 14 of which have been reclassified to the following genera: *Blautia, Mediterraneibacter, Trichococcus,* or *Hominimerdicola*. Of the remaining 18 genomes, nine overlapped with the 124 NCBI genomes and nine had no significant scaffoldin or DocGH detected by IPS or AF2. To comprehensively identify cohesins by primary sequence analysis, we combined BLAST and InterProScan (v. 5.59-91.0) searches. InterProScan was used to detect cohesins matching sequence profiles defined in the Conserved Domain Database (CDD) and Pfam: type-I (cd08548), type-II (cd08547), type-III (cd08759), and general cohesin (PF00963). Additionally, dockerin domains were detected using CDD profiles (type-I (cd14256), type-II (cd14254), type-III (cd14255)) to identify those genomes with greater than 15 dockerin-containing proteins. In parallel, BLAST analyses were performed using select representative cohesin domains from each type as queries against each genome, applying an E-value cutoff of 1e-4.^17,46^ These methods revealed 361 cohesin domains across 33 *Ruminococcus* genomes that have greater than 15 dockerin proteins.

Full proteome structures were predicted using AF2 for 11 *Ruminococcus* species for which high-quality genome sequence data is available. For the remaining 22 metagenome-assembled genomes, only proteins containing a signal peptide by SignalP (v. 6.0)^47^ were modeled because cellulosome-associated proteins are expected to be secreted. This greatly reduced the computational time required for AlphaFold predictions. Proteomes were subsequently broken down into individual domains by the Unified Domain Cutter (UniDoc)^25^ yielding 94,234 domains. Next, domains were structurally predicted by AlphaFold2. For each sequence, three models were generated, and the model with the highest pLDDT score was selected for structural analysis. Predicted models were structurally compared to template type-1,-2, and –3 cohesin structures (type-I (PDB:1OHZ), type-II (PDB:2BM3) and type-III (PDB:2ZF9) cohesins) using TMalign.^26^ A carbohydrate binding module family 3 (CBM3) protein structure (PDB:6UFW) was used as a negative control since it displays similar folds to cohesin domains. A structural comparison with a TM-score above 0.65 was considered a quality match to the reference models. Overall, there were 225 matches to type-I, 13 matches to type-II, and 603 matches to type-III. If a domain had a TM-score higher than 0.65 to multiple reference models, the highest scoring reference model was taken. In addition to the 361 BLAST or InterProScan identified cohesin domains, our structural comparison analysis revealed 177 more cohesin domains with a TM-score above 0.65.

A structure-based tree was constructed using TM-scores from an all-on-all comparison of the AlphaFold-only identified cohesin domains along with PDB annotated cohesin domains. A distance matrix was calculated from TM-score values (1-TM-score) and used to build a tree with MEGA12.^48^ Given the relatively tight range of acceptable TM-scores that signify structural significance, 0.65-1.00, it is difficult to define clear grouping boundaries within the structural tree. To this point, groups were established by creating the largest cluster of domains from a common node that consisted of at least nine members with average TM-score similarity amongst themselves greater than 0.80.

Further, a sequence-based tree was constructed using the amino acid sequences of the cohesin domains described above. A multiple sequence alignment (MSA) was generated by the <u>MU</u>ltiple <u>S</u>equence <u>C</u>omparison by <u>L</u>og-<u>E</u>xpectation (MUSCLE) program.^49^ For visualization, a sequence similarity network (SNN) was generated using the MUSCLA MSA with Cytoscape.^50^ An alignment threshold score of 30% was set for clustering parameters.

### Structural determination of *R. callidus* cohesin domains

Expression constructs for the *R. callidus* VPI 57-31 (ATCC 27760) putative cohesins, including an N-terminal TEV-cleavable His6 tag, were synthesized and assembled into pET29b (Twist Bioscience). Plasmids were transformed into *E*. *coli* BL21-Gold (DE3) and expressed using Overnight Express Instant TB media (Novagen) for 3 days at 25°C. Cells were harvested by centrifugation, resuspended in Buffer A (50 Tris pH 8, 300 NaCl, 20 mM 300 imidazole) supplemented with EDTA (1 mM) and PMSF (1 mM), and disrupted using an Emulsiflex C-3 high pressure homogenizer (Avestin). The lysate was clarified by centrifugation and the supernatant was loaded on a 5 ml HisTrap Crude FF column (Cytiva) equilibrated in Buffer A. Bound protein was eluted with Buffer B (Buffer A with 300 mM Imidazole) directly onto a Superdex 75 column (Cytiva) equilibrated in TBS (20 mM Tris pH 7.5, 150 mM NaCl). Fractions were analyzed by SDS-PAGE and those containing the protein of interest were concentrated to 10-75 mg/ml for crystallization screening. The His6 tag was cleaved from target proteins that failed to yield crystallization leads and were subjected to further screening. Targets that continued to prove recalcitrant to crystallization were dialyzed into 25 mM HEPES pH 7.6, 150 mM NaCl and solvent accessible lysines were methylated following the protocol of Kim et al..^51^ The methylated target proteins were subsequently subjected to crystallization screening.

### Genome similarity and phylogenetic reconstruction using average nucleotide identity

Whole-genome relatedness among *Ruminococcus* isolates was evaluated using Average Nucleotide Identity (ANI). Pairwise ANI values were calculated using the ANIb algorithm implemented in the pyani-plus software package,^52^ which performs BLAST-based^53^ homologous sequence comparisons and quantifies nucleotide identity across conserved genomic segments. The resulting values were exported as a square matrix, representing all pairwise genome comparisons.

To prepare the data for phylogenetic inference, genome identifiers were standardized to remove file-format artifacts and ensure consistent taxon labeling. Pairwise genomic distance values were subsequently derived from the identity matrix using D = 1 – ANI, where ANI values were treated as proportional measures of nucleotide similarity. Phylogenetic trees were reconstructed using two classical distance-based clustering methods, Neighbor-Joining (NJ)^54^ and Unweighted Pair Group Method with Arithmetic Mean (UPGMA), enabling assessment of genome clustering under different evolutionary assumptions. NJ was selected to minimize total branch length under a minimum-evolution framework, whereas UPGMA applied hierarchical clustering based on average linkage. Negative or undefined branch lengths were corrected to avoid distortions in topology.

## Acknowledgments

This work used computational and storage services associated with the Hoffman2 Cluster which is operated by the UCLA Office of Advanced Research Computing’s Research Technology Group. We would also like to acknowledge and thank the computational resources of the UCLA DOE-IGP cluster.

## Author Contributions

R.T.C. designed research and wrote the paper; C.M. and A.T. designed research, performed experiments, analyzed data, and wrote the paper; M.A. and S.M.H. contributed new reagents and analytical tools; R.P.G. performed experiments and analyzed data; M.P. contributed new reagents and analytical tools; M.R.S. performed experiments and analyzed data.

## Classification

Biological Sciences/Microbiology

## Figures and Tables

**Figure S1.**
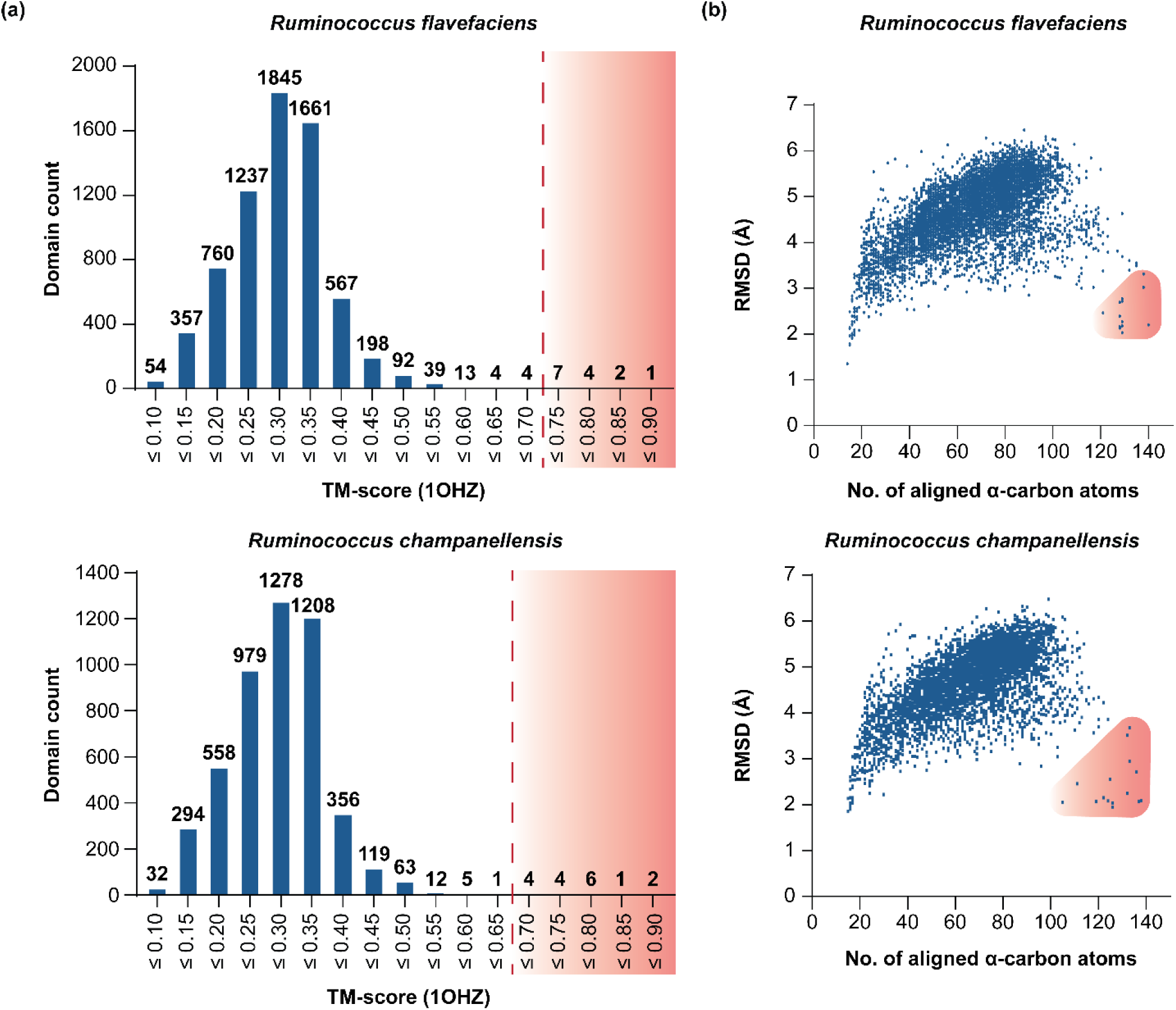
Established TM-score threshold is consistent across other cellulosomal *Ruminococcus*. **(A)** Representative plots showing TM-score versus domain count and **(B)** RMSD α-carbon coordinate difference versus number of aligned α-carbon atoms for predicted structures in *Ruminococcus flavefaciens* (top) and *R. champanellensis* (bottom). 6845 and 4922 individual domains were compared to a type I cohesin (PDB: 1OHZ) for *R. flavefaciens* and *R. champanellensis*, respectively. The threshold TM-score used for cohesin domain identification is shown as a dashed red line that separates a large low-scoring population of unrelated structures (left) from a high-scoring structurally related population (right).

**Figure S2.**
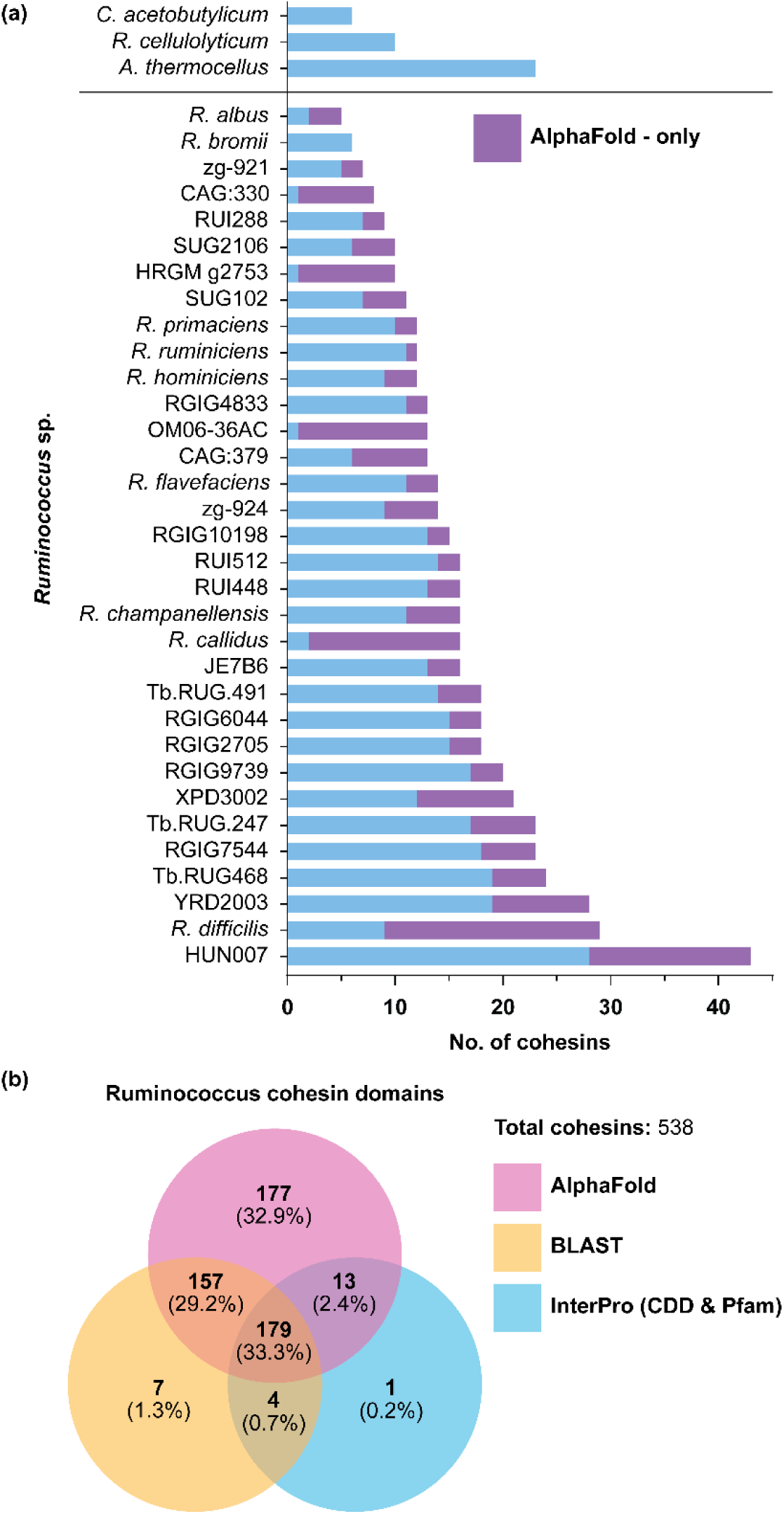
AlphaFold identifies a large number of novel cohesin domains in *Ruminococcus* proteomes. **(A)** Bar graph showing the total number of cohesin domains identified per *Ruminococcus* proteome. Total bar height represents the number of cohesin domains identified across all annotation methods used (AF2, InterProScan, BLAST). Segments colored purple indicate AF2-only detected cohesins. Also shown are counts for representative non-*Ruminococcus* cellulosome producing bacteria from the *Acetivibrio* (*Acetivibrio thermocellus*), *Ruminoclostridium* (*Ruminoclostridium cellulolyticum*), and *Clostridium* (*Clostridium acetobutylicum*) genera. **(B)** Venn diagram of the total number of cohesin domains identified by AF2 (pink), InterProScan (cyan), and BLAST (yellow) in analyzed *Ruminococcus* proteomes.

**Figure S3.**
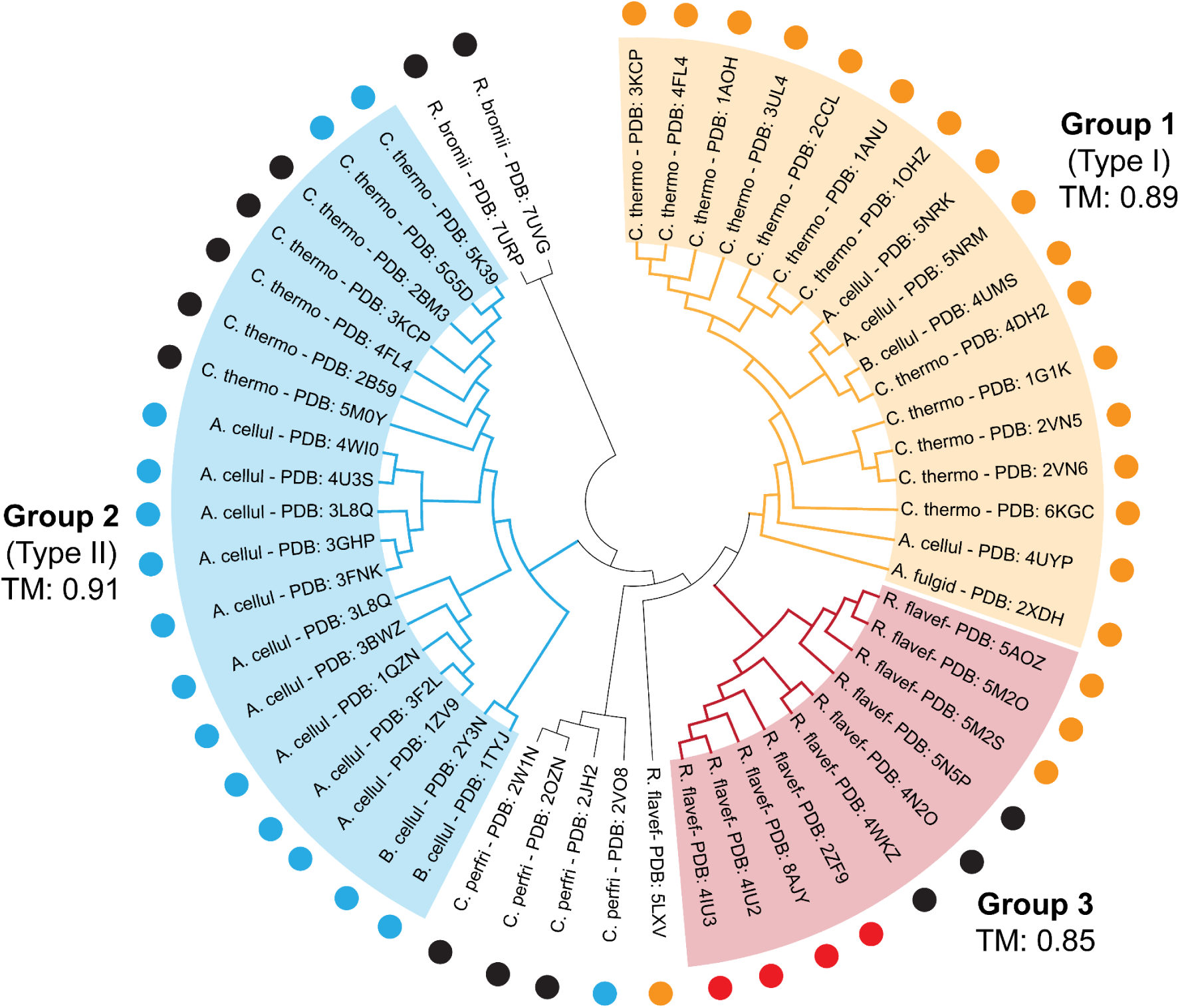
Structure-based dendrogram of experimentally determined cohesin structures. Structural dendrogram of all 52 experimentally determined cohesin atomic structures deposited in the PDB. The dendrogram was constructed from an all-on-all TM-score matrix. The majority of structures can be classified into structural groups 1 to 3 whose members are frequently assigned to type I to III cohesin families based on their primary sequences, respectively. In the figure these distinct primary sequences are indicted with dots (type I (yellow, cd08548), type II (cyan, cd08547), type III (red, cd08759), and un-typed (black, PF00963). Additional, sparsely populated structural families are also formed by cohesins from *R. bromii*, *C. perfringens*, and *R. flavefaciens* species. The minimum TM-score for inclusion into groups 1 to 3 is indicated.

**Figure S4.**
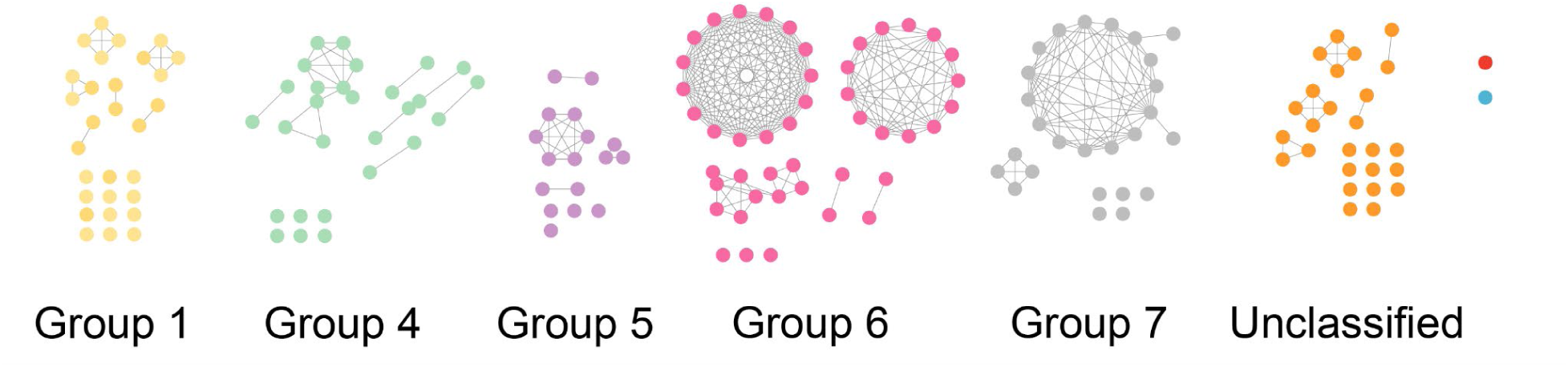
Sequence Similarity Network (SSN) of 177 AF2-only identified ruminococcal cohesins. Sequence similarity network (SSN) of 177 AF2-only identified cohesins was created by EFI-EST and visualized with CytoScape. Three additional sequences are included as representative type-1, –2, and –3 cohesins from the PDB (type-1: 1OHZ, type-2: 1TYJ, type-3: 2ZF9). A sequence similarity cutoff of 30% was used to generate clusters. Colors of nodes denote the structural group each cohesin domain belongs to; Yellow: Group1, Green: Group 4, Purple: Group 5, Pink: Group 6, Grey: Group7, Orange: No assigned group.

**Figure S5.**
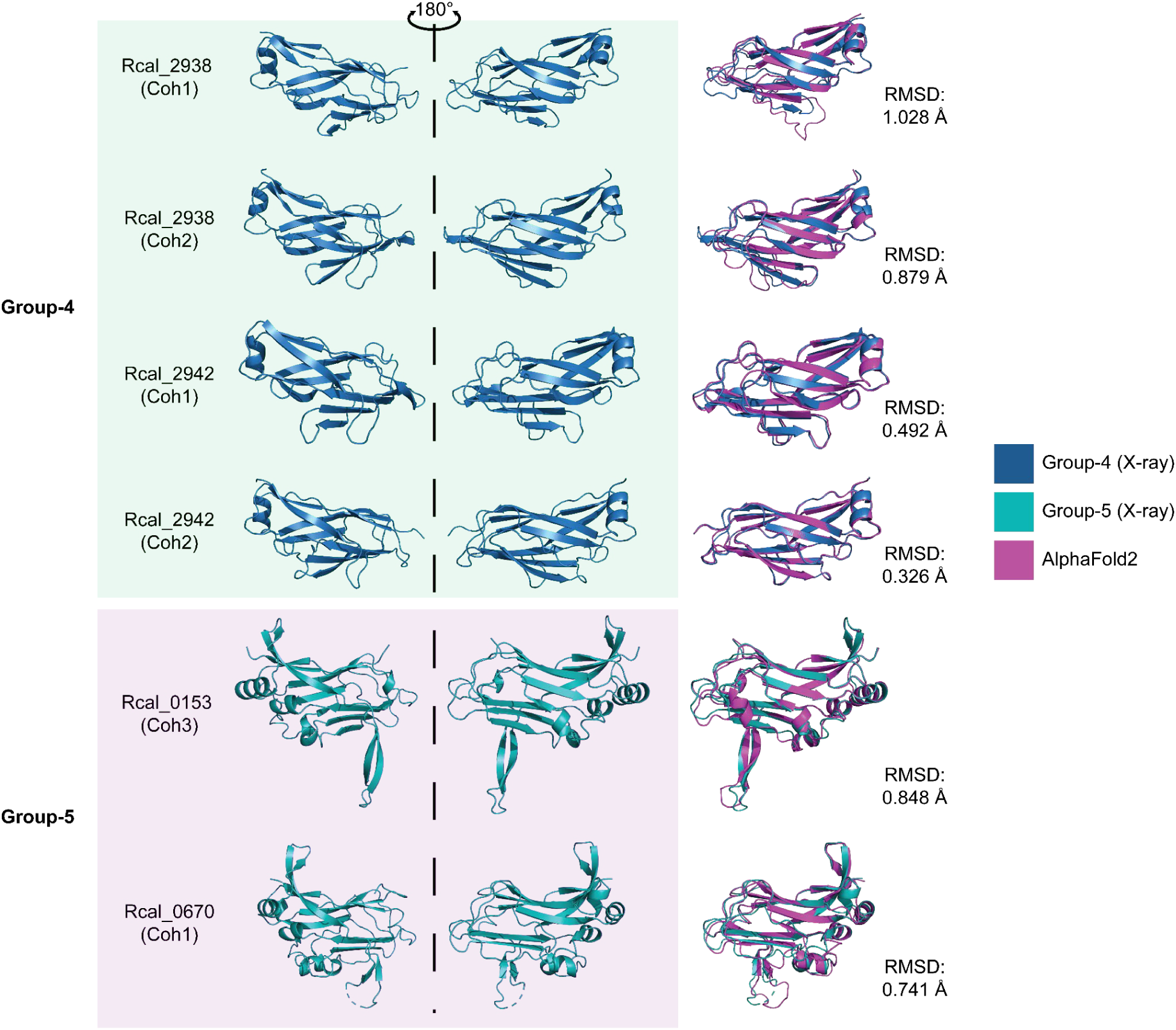
Crystal structures of novel *R. callidus* cohesins. Cartoon models of experimentally determined structures for six additional putative cohesins in *R. callidus* scaffoldin-like proteins. Overlays of the crystal structure shown here and the corresponding predicted AlphaFold2 (purple) model are shown on the right with the calculated backbone root mean square deviation (RMSD) between the two structures.

**Figure S6.**
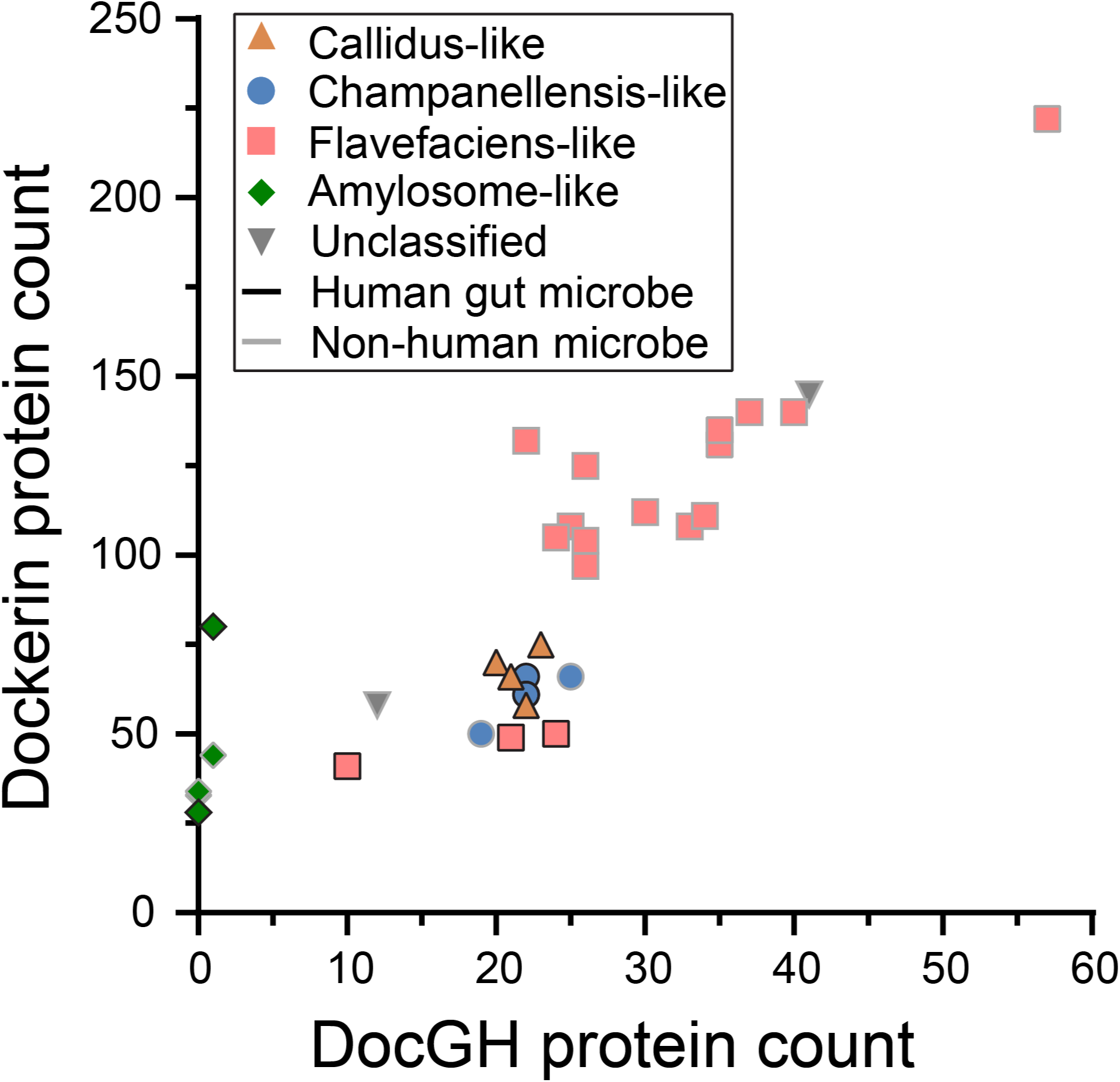
Cellulolytic capacity of ruminococcal species. Counts for DocGH and Dockerin-containing proteins are plotted for species within each cellulosome-archetype. Those species outlined in black represent human-gut associated microbes and those in grey are non-human gut associated. The unclassified species are *Ruminococcus albus* (58, 12) and *Ruminococcus HUN007* (57, 222) which has a unique cellulosome architecture.

**Figure S7.**
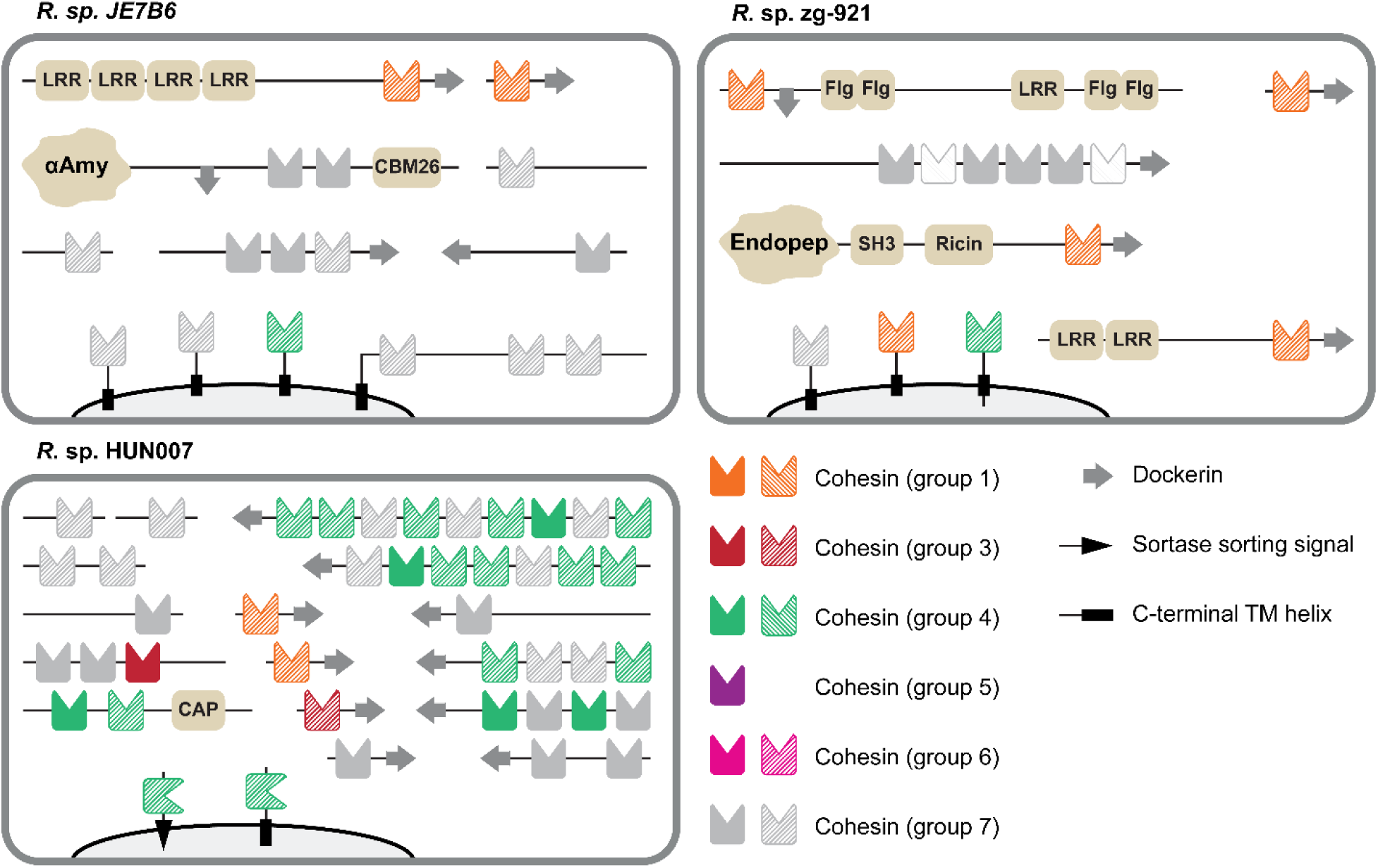
Cellulosome-like architectures in non-human associated Ruminococci. Cartoon schematic of the cellulosomal proteins in two species analyzed that did not follow the archetypes of other *Ruminococcus* cellulosomes established in Fig. 5, *Ruminococcus bromii* and *Ruminococcus* sp. HUN007. Color and shading for each cohesin domain follows the same convention established in Fig. 5. Along with those represented in Fig. 5, these cartoon schematics represent all cellulosome-like assemblies discussed, with the exception of *Ruminococcus albus*. Additional domains within the scaffoldin and scaffoldin-like proteins are also indicated: **LRR** – Leucine Rich Repeat, **αAmy** – α-Amylase, **CBM26** – Carbohydrate Binding Module Family 26, **Flg** – Listeria-Bacteroides repeat domain, **Endopep** – endopeptidase domain, **SH3** – SRC Homology 3 domain, **Ricin** – Ricin B-like lectin, **X** – X2 domain, **GH** – glycoside hydrolase, **CAP** – Cysteine-rich secretory / Antigen 5 / Pathogenesis-related 1 domain.

**Table S1:**
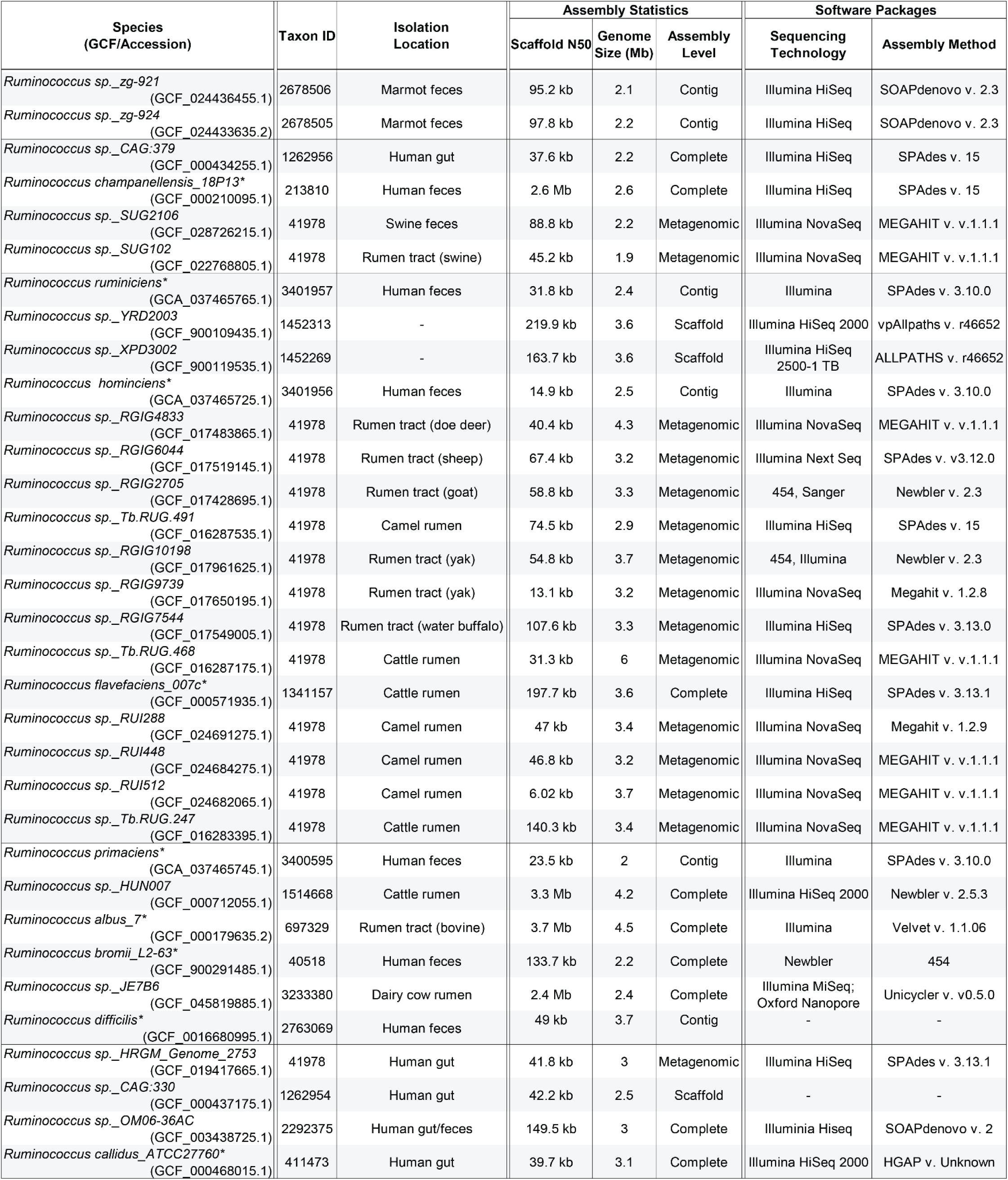
*Ruminococcus spp.* Genome metrics. List of the genome quality statistics for all analyzed *Ruminococcus* genomes including species name, GCF reference, taxonomy ID, isolation location, assembly statistics, and associated software packages used to analyze the genome. The N50 contig value is the shortest length contig that when added with all other contigs greater in length sums to roughly 50% of the total assembly length. All analyzed *Ruminococcus* genomes were annotated using Prokka.^50^ A * symbol indicates *Ruminococcus* species listed on the List of Prokaryotic Names with Standing in Nomenclature. In our analysis, we also inspected the genomes of several species that were originally assigned as *Ruminococcus* bacteria, but have since been reassigned to *Blautia*, *Mediterraneibacter*, *Trichococcus*, and *Hominimerdicola* genera. The additional nine genomes that were analyzed were not found to have cellulosome-like structures after AF analysis: *R. sp._RGIG8444, R. intestinalis, R. sp. BSD2780120874_150323_B10, R. equi, R. sp. FMB-CY1, R. sp. JL13D9, R. sp. YE71, R. sp. YE78, R. bovis, R. sp. FC2018*.

**Table S2.** X-ray data-collection and refinement statistics for *R. callidus* cohesins: *Values in parentheses are for the highest resolution shell.

|  | ScaB<br>(Rcal_0153) |  | Rcal_0670 | Rcal_2938 |  | Rcal_2942 |  |  |
| --- | --- | --- | --- | --- | --- | --- | --- | --- |
| Cohesin No.<br>Structural Group | Coh3<br>5 | Coh4<br>5 | Coh1<br>5 | Coh1<br>4 | Coh2<br>4 | Coh1<br>4 | Coh2<br>4 | Coh4<br>4 |
| **Data Collection** |  |  |  |  |  |  |  |  |
| Beamline | APS 24-ID-E | APS 24-ID-E | APS 24-ID-E | APS 24-ID-E | APS 24-ID-E | APS 24-ID-E | APS 24-ID-E | APS 24-ID-E |
| Space Group | P2<sub>1</sub> | C222<sub>1</sub> | P2<sub>1</sub>2<sub>1</sub>2<sub>1</sub> | P4<sub>1</sub> | C222<sub>1</sub> | P2<sub>1</sub>2<sub>1</sub>2<sub>1</sub> | P2<sub>1</sub> | P2<sub>1</sub>2<sub>1</sub>2<sub>1</sub> |
| Resolution (Å) | 2.80 (2.87-2.80)* | 1.80 (1.85-1.80)* | 2.10 (2.16-2.10)* | 1.60 (1.64-1.60)* | 2.00 (2.05-2.00)* | 1.90 (1.95-1.90)* | 1.90 (1.95-1.90)* | 1.20 (1.27-1.20)* |
| Unit cell lengths: a,b,c (Å) | 92.2, 172.3, 92.3 | 42.5, 84.6, 118.4 | 42.6, 44.0, 267.3 | 52.3, 52.3, 101.8 | 86.8, 88.9, 82.3 | 121.9, 121.5, 60.3 | 40.0, 147.5, 61.2 | 34.0, 37.2, 110.4 |
| Unit cell angles: α,β,γ (°) | 90, 106.2, 90 | 90, 90, 90 | 90, 90, 90 | 90, 90, 90 | 90, 90, 90 | 90, 90, 90 | 90, 96.1, 90 | 90, 90, 90 |
| Measured reflections | 390062 (29927) | 157327 (5854) | 311825 (20916) | 753780 (53127) | 140382 (9962) | 349805 (23819) | 314927 (14870) | 565095 (85641) |
| Unique reflections | 67393 (4981) | 19021 (976) | 30304 (2151) | 36129 (2684) | 21748 (1563) | 70378 (5039) | 53864 (3896) | 84166 (13514) |
| Overall completeness (%) | 99.4 (99.1) | 94.1 (65.5) | 99.9 (99.5) | 100.0 (99.5) | 99.5 (99.5) | 98.8 (97.1) | 97.3 (82.6) | 99.4 (98.5) |
| Overall redundancy | 5.8 (6.0) | 8.3 (6.0) | 10.2 (9.7) | 20.9 (19.8) | 6.5 (6.4) | 5.0 (4.7) | 5.8 (3.8) | 6.7 (6.3) |
| Overall R<sub>merge</sub> | 0.201 (1.747) | 0.128 (0.591) | 0.210 (1.33) | 0.140 (1.53) | 0.082 (0.929) | 0.108 (0.87) | 0.108 (0.785) | 0.095 (1.03) |
| CC<sub>1/2</sub> | 98.8 (27.7) | 99.1 (76.8) | 99.5 (53.5) | 99.9 (68.2) | 99.8 (64.1) | 99.4 (66.0) | 99.7 (69.4) | 99.8 (55.3) |
| Overall I/σ | 8.5 (1.6) | 11.6 (2.6) | 9.8 (2.0) | 13.9 (2.8) | 14.3 (2.5) | 9.7 (2.3) | 11.2 (1.9) | 10.3 (1.6) |
| **Refinement** |  |  |  |  |  |  |  |  |
| R<sub>work</sub> / R<sub>free</sub> | 0.186 / 0.202 | 0.188 / 0.234 | 0.202 / 0.231 | 0.165 / 0.198 | 0.176 / 0.212 | 0.169 / 0.202 | 0.169 / 0.202 | 0.148 / 0.174 |
| RMSD bond length (Å) | 0.007 | 0.006 | 0.008 | 0.017 | 0.007 | 0.008 | 0.01 | 0.005 |
| RMSD angle (°) | 1.3 | 1.2 | 0.9 | 2 | 0.8 | 1.5 | 1.5 | 0.8 |
| No. of protein atoms | 17934 | 1754 | 3258 | 2155 | 2111 | 4769 | 4432 | 2577 |
| No. of water atoms | 28 | 184 | 135 | 160 | 119 | 364 | 270 | 145 |
| No. of other solvent atoms | 65 | 4 | 0 | 17 | 0 | 79 | 58 | 0 |
| Avg. B-factor of protein (Å<sup>2</sup>) | 64.9 | 18.6 | 34.2 | 27.2 | 40.5 | 35.4 | 36.8 | 17.1 |
| Avg. B-factor of water (Å<sup>2</sup>) | 36.2 | 23.6 | 32 | 32.6 | 43 | 38.2 | 36.7 | 25.1 |
| Avg. B-factor of other solvent (Å<sup>2</sup>) | 64.5 | 33.8 | N/A | 45.6 | N/A | 53.5 | 55.8 | N/A |
| PDB ID code | #### | #### | #### | #### | #### | #### | #### | #### |

**Table S3.**
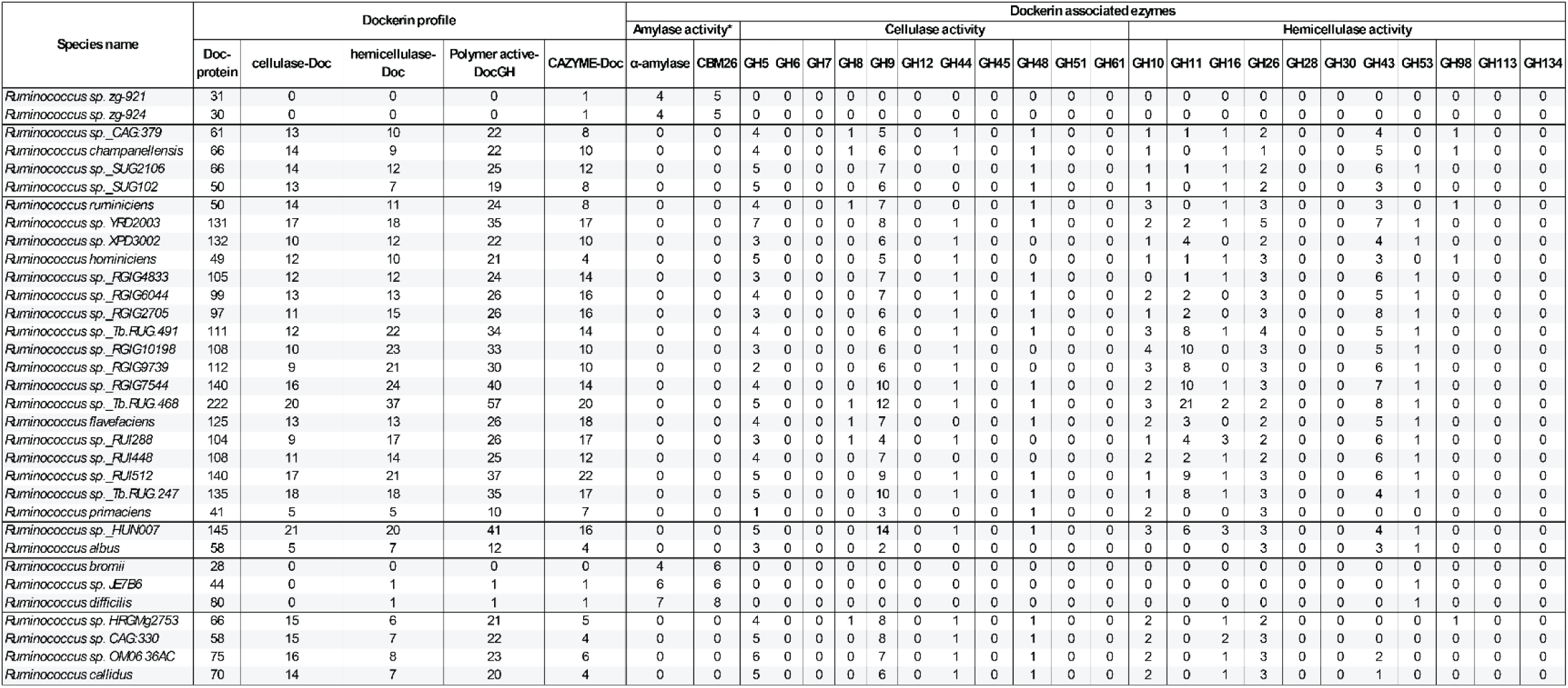
Predicted enzyme content of cellulosome displaying bacteria: Dockerin-enzyme content of 33 *Ruminococcus* species grouped together within their phylogenetic relationships. Total dockerin proteins, cellulase-doc, hemicellulase-doc, polymer-active DocGH (combined cellulase and hemicellulase-DocGH content), and CAZyme-Doc proteins are noted. The dockerin-enzyme profile is divided into protein counts of dockerin-fused amylase, cellulase, and hemicellulase active enzymes. *CBM26 is a starch binding carbohydrate binding module present in *Ruminococcus* species that contain Amylase-Doc proteins.

**Table S4.** Putative Scaffoldin proteins and cohesin domains identified in *Ruminococcus callidus*: Tabulation of putative cohesin domain-containing scaffoldin proteins for *R. callidus* including; protein accession code (Protein ID), Gene Locus, cohesin structural group (Group), TM-align scores to each representative cohesin domain (AF2 / TM-align), InterProScan sequence identifier annotation (CDD and Pfam), if the domain was identified by BLAST (BLAST Identified), and domain arrangement according to structurally and sequence-identified domains (Modular Arrangement). Abbreviations: cohesin (**Coh**), dockerin (**Doc**), cysteine-rich secretory / antigen 5 / pathogenesis-related 1 domain (**CAP**), sortase sorting signal (**LPXTG**), domain of unknown function (**DUF5620**), signal peptide (**SIGN**), X-module (**X**).

| Protein ID | Gene locus | Group | AF2 / TM-align |  |  | InterProScan |  | BLAST identified | Modular arrangement |
| --- | --- | --- | --- | --- | --- | --- | --- | --- | --- |
|  |  |  | Type I (1OHZ) | Type II (1TYJ) | Type III (2ZF9) | CDD | Pfam |  |  |
| WP_021681153 | Rcal_0005 (ScaE-like) | 4 | 0.716 | 0.620 | 0.618 | - | - | - | SIGN-Coh-LPXTG |
| WP_021682353 | Rcal_0152 (ScaC-like) | 1 | 0.870 | 0.708 | 0.686 | cd08548 | PF00963 | ✓ | SIGN-Coh-Doc |
| WP_370809552 | Rcal_0153 (ScaB-like) | 6 | 0.718 | 0.674 | 0.644 | - | - | - | SIGN-Coh-Coh-Coh-Coh-X-Doc |
|  |  | 5 | 0.798 | 0.709 | 0.686 | - | - | - |  |
|  |  | 5 | 0.795 | 0.711 | 0.685 | - | - | - |  |
|  |  | 5 | 0.794 | 0.692 | 0.655 | - | - | - |  |
| WP_051343558 | Rcal_0670 | 5 | 0.765 | 0.700 | 0.694 | - | - | - | SIGN-DUF5620--Coh-Coh-- |
|  |  | 5 | 0.763 | 0.688 | 0.650 | - | - | - |  |
| WP_021682326 | Rcal_1096 | 7 | 0.544 | 0.448 | 0.411 | - | PF00963 | ✓ | SIGN-Coh |
| WP_021680365 | Rcal_2513 | 6 | 0.738 | 0.671 | 0.634 | - | - | - | SIGN-Coh-- |
| WP_169725972 | Rcal_2938 | 4 | 0.706 | 0.596 | 0.602 | - | - | - | SIGN-CAP-Coh-Coh-Doc |
|  |  | 4 | 0.717 | 0.610 | 0.623 | - | - | - |  |
| WP_021680502 | Rcal_2942 | 4 | 0.764 | 0.643 | 0.643 | - | - | - | SIGN-Coh-Coh-Coh-Coh-Doc |
|  |  | 4 | 0.762 | 0.635 | 0.634 | - | - | - |  |
|  |  | 4 | 0.775 | 0.663 | 0.663 | - | - | - |  |
|  |  | 4 | 0.777 | 0.649 | 0.639 | - | - | - |  |

**Table S5.**
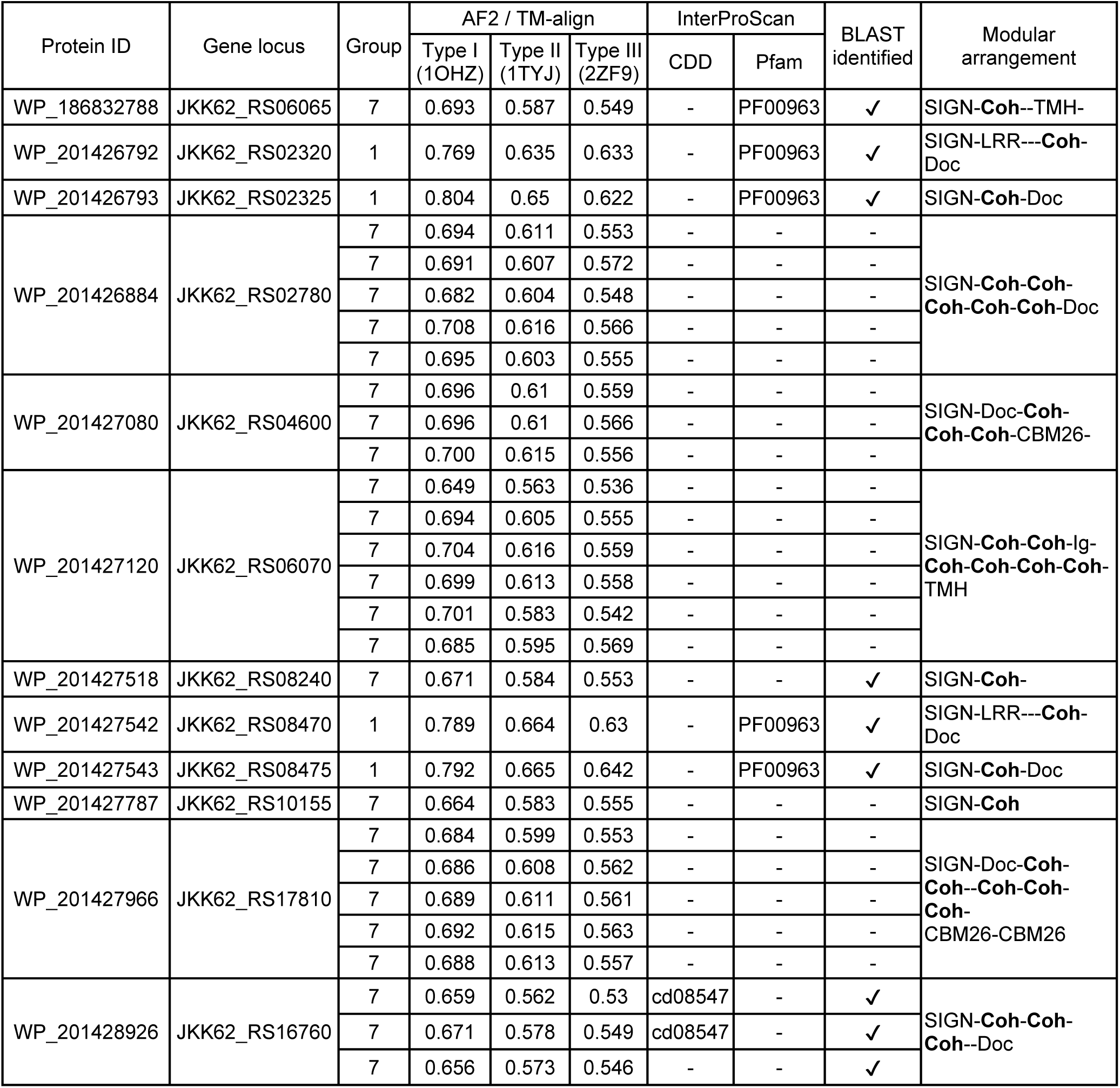
Putative Scaffoldin proteins and cohesin domains identified in *Ruminococcus difficilis*: Tabulation of putative cohesin domain-containing scaffoldin proteins for *R. difficilis* including; protein accession code (Protein ID), Gene Locus, cohesin structural group (Group), TM-align scores to each representative cohesin domain (AF2 / TM-align), InterProScan sequence identifier annotation (CDD and Pfam), if the domain was identified by BLAST (BLAST Identified), and domain arrangement according to structurally and sequence-identified domains (Modular Arrangement). Abbreviations: cohesin (**Coh**), dockerin (**Doc**), carbohydrate binding module, family 26 (**CBM26**), predicted transmembrane helix region (**TMH**), leucine-rich repeat region (**LRR**), signal peptide (**SIGN**)

